# CaMKIIa+ neurons in the bed nucleus of the stria terminalis modulate pace of natural reward seeking depending on internal state

**DOI:** 10.1101/2023.09.17.558107

**Authors:** Patty T. Huijgens, Roy Heijkoop, Louk J.M.J. Vanderschuren, Heidi M.B. Lesscher, Eelke M.S. Snoeren

## Abstract

This study aims to investigate the underlying neurobiological mechanisms that regulate natural reward seeking behaviors, specifically in the context of sexual behavior and sucrose self-administration. The role of CaMKIIa+ neurons in the bed nucleus of the stria terminalis (BNST) was explored using chemogenetic silencing and -stimulation. Additionally, the study examined how these effects interacted with the internal state of the animals. Through detailed behavioral analysis, it was demonstrated that CaMKIIa+ neurons in the BNST play a significant role in the regulation of both sexual behavior and sucrose self-administration. Although the behavioral outcome measures differed between the two behaviors, the regulatory role of the CaMKIIa+ neurons in the BNST was found to converge on the modulation of the pacing of engagement in these behaviors in male rats. Moreover, our study confirmed that the internal physiological state of the animal affects how the BNST modulates these behaviors. These findings suggest that different types of natural rewards may recruit a similar brain circuitry to regulate the display of motivated behaviors. Overall, this research provides valuable insights into the neural mechanisms underlying natural reward seeking and sheds light on the interconnected nature of reward-related behaviors in male rats.

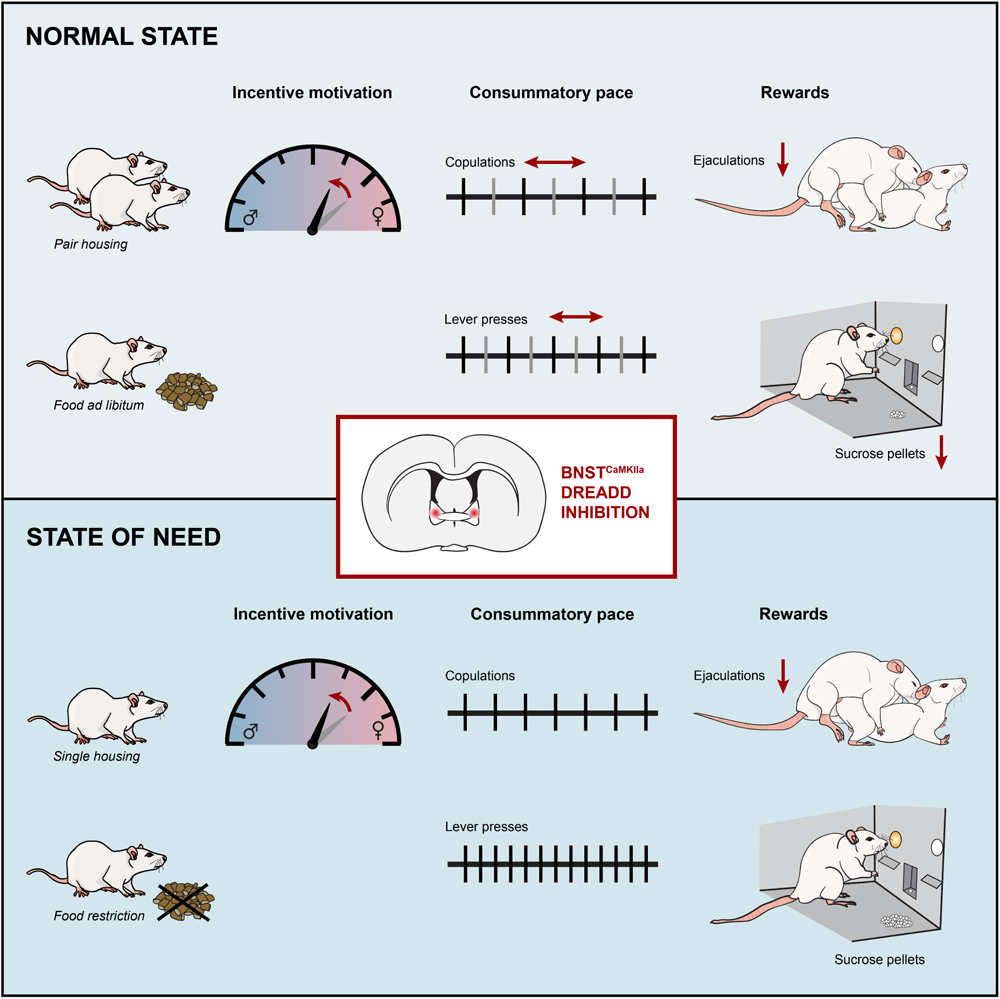

## Introduction

Different types of stimuli can have attractive and motivational values that induce approach and consummatory behavior. Food, social interactions, and sexual partners are examples of ‘natural rewards’, because they induce behavior that is needed to survive, or to ensure survival of the species. At the same time, they are examples of ‘intrinsic rewards’, rewards that have an unconditioned, inherently pleasurable value. Shared across the individual’s engagement with each naturally rewarding stimulus are specific behavioral states that can be described as motivation, approach, consumption, and satiety. In addition, consummation of natural rewards requires an incentive motivation to engage in the behavior, and a motivation to continue the behavior after it has commenced. These motivational states can differ and can be assessed separately using extensive behavioral analysis. Sexual behavior and sucrose consummation are innately motivated and rewarding behaviors. In experienced male rats, both ejaculation and intromissions are rewarding, and can induce conditioned place preference, with ejaculation having a greater reward value than intromissions (Camacho et al. 2009; Tenk et al. 2009). Sucrose consumption also induces robust conditioned place preference in male rats (Patel et al. 2020; Ågmo et al. 1995). In both sexual behavior and sucrose consumption a level of motivation for an incentive stimulus is followed by a behavioral response of approach and consumption. Natural reward seeking behaviors are usually studied separately, and it is not fully known whether specific micropatterns of the behavior overlap and to what extent the underlying neurobiological mechanisms converge. Nonetheless, given that every naturally rewarding stimulus induces a similar sequence of behavioral states that culminates in consumption and subsequent satiety, it is plausible that various natural rewards may elicit shared behavioral responses, which are regulated by common neural mechanisms.

The bed nucleus of the stria terminalis (BNST) is implicated in stress, anxiety, drug seeking, addiction, and socio-sexual behavior (reviewed in: (Flanigan and Kash 2022; Lebow and Chen 2016)). Through its projection to the ventral tegmental area, the BNST modulates reward and aversion (Jennings et al. 2013b; Soden et al. 2022), while connections with the amygdala, medial prefrontal cortex, and hypothalamus are involved in processing mood and arousal states, and monitoring of food and fluid intake (Davis et al. 2010; Dong et al. 2001; Dong and Swanson 2004; Hammack et al. 2009; Radley and Sawchenko 2011; Rodriguez-Romaguera et al. 2020). Consequently, it has been proposed that the BNST has a function in valence surveillance, meaning that it weighs various types of information such as environmental context, historical experience and internal state, in order to guide an appropriate response towards a stimulus (Lebow and Chen 2016).

The BNST has also been shown to have a role in sexual behavior. In male rats, c-Fos+ cell density in the posteromedial division of the BNST is increased upon chemosensory investigation of a female as well as upon copulation, especially after reaching an ejaculation (Coolen et al. 1996; 1997; 1998). Lesions of the BNST result in an increased ejaculation latency, a higher number of mounts and intromissions preceding ejaculation, and both longer interintromission and post-ejaculatory intervals (Emery and Sachs 1976; Valcourt and Sachs 1979). The BNST is a highly heterogenous brain region, in which as many as 18 subnuclei (Lebow and Chen 2016), and 41 transcriptionally distinct neuron populations have been identified in mice (Welch et al. 2019). What the role of the BNST is in sexual incentive motivation and which cells in particular play a role in sexual behavior in rats remains unknown. In mice, it was shown that at least aromatase+, cholecystokinin+ and Esr2+ neurons are involved in preference for female pheromones (Bayless et al. 2019a; Giardino et al. 2018) and the regulation of copulatory behaviors (Bayless et al. 2019a; Zhou et al. 2023). Excitotoxic lesions of the BNST has also been shown to eliminate sexual odor preference in sexually naïve hamster males (Been and Petrulis 2010). In mice, 15% of all neurons in the anterodorsal BNST and 8.6% in the ventral BNST express CaMKII (Nguyen et al. 2016). We have previously demonstrated that CaMKIIa+ neurons in the medial amygdala modulate ejaculation latency through regulation of number of intromissions preceding ejaculation. The role of BNST^CaMKII^ neurons in the BNST has not yet been studied in natural reward seeking behaviors.

Many studies have investigated the role of the BNST in self-administration of drugs of abuse, drug seeking, and relapse. However, self-administration of more natural consumable rewards, such as sucrose, has not been studied as extensively. It is known, though, that intra-oral sucrose infusions increase dopamine release in the BNST (Park et al. 2012), and dopamine antagonist injections into the BNST reduce responding to sucrose in a binge eating paradigm (Maracle et al. 2019). Moreover, sucrose-seeking under fixed ratio (FR) paradigms, but not under progressive ratio (PR), increases c-Fos+ cell density in the anterior and posterior divisions of the BNST (Figlewicz et al. 2011). So far, only a few studies have been performed in the BNST to explore the functional role of the BNST in sucrose consumption. It has been shown that GABA knockdown in CRF-expressing BNST neurons reduces motivation to work for sucrose, an effect that was selective for male mice (Gianessi et al. 2023). In addition, while stimulation of serotonin 5HT(2c) receptor containing cells in the BNST reduces sucrose intake in only female mice (Flanigan et al. 2023b), inhibition of these neurons does not seem to have an effect in both sexes (Flanigan et al. 2023a).

Although the involvement of the BNST has been shown in both sexual behavior and sucrose consumption, the precise characteristics of its role in regulating the specific behavioral states and micropatterns remain unexplored. Furthermore, the question remains whether these two natural rewards may elicit shared behavioral responses, and if so, whether these responses are regulated by common neural mechanisms. Therefore, in this study, the role of CaMKIIa+ neurons in the BNST was explored using chemogenetic silencing and -stimulation. Through detailed behavioral analysis, we demonstrate that CaMKIIa+ neurons in the BNST play a significant role in the regulation of both sexual behavior and sucrose intake. Although the behavioral outcome measures differed between these two behaviors, the regulatory role of the BNST was found to converge on the modulation of the pacing of engagement in these behaviors in male rats. Moreover, these effects interacted with behavior-specific internal states. These findings imply a shared neural circuitry for regulating motivated behaviors across various natural rewards.

## Results

The first aim of the study was to explore the role of BNST^CaMKIIa^ neurons in male rat sexual behavior. Thirty-six male Wistar rats were first sexually trained in the copulation test set-up, and then divided into three groups (n=12) with homologous phenotypes regarding number of ejaculations reached in 30 minutes. Then, CaMKIIa+ neurons in the BNST were targeted by stereotaxically injecting an AAV encoding either inhibitory (Gi), stimulatory (Gq) or control (Sham) DREADDs under the CaMKIIa promoter (Fig. 1A).

**Figure 1.**
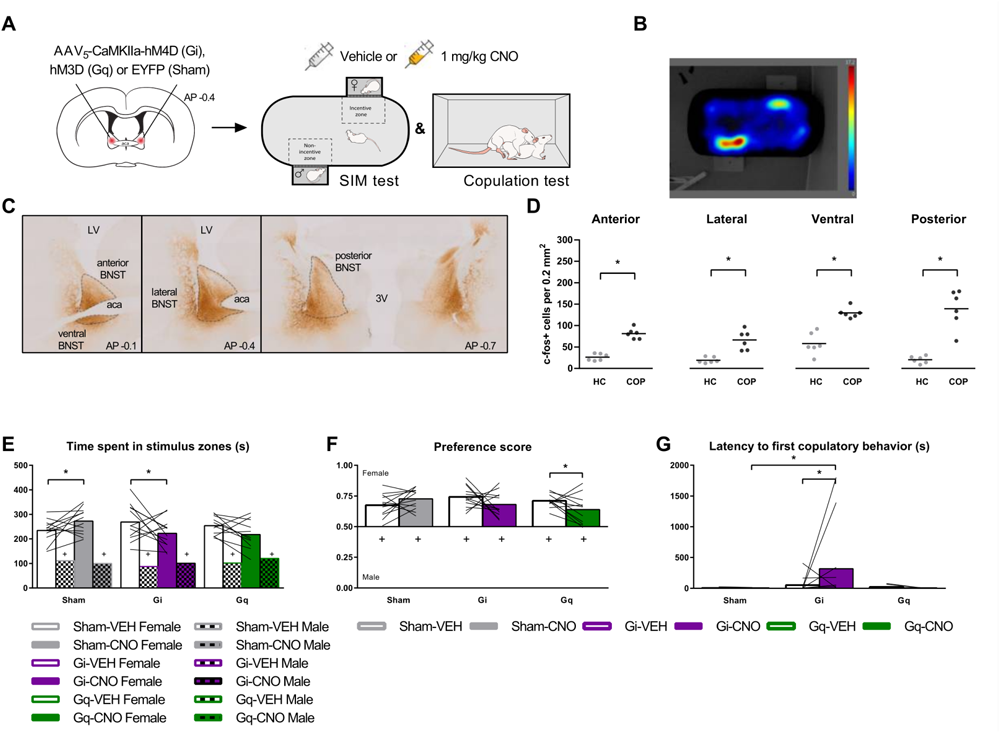
Effects of DREADDs on sexual incentive motivation. **(A)** Bilateral viral targeting of the bed nucleus of the stria terminalis (BNST), and experimental timeline. Subject males were tested in the sexual incentive motivation (SIM) and copulation test after vehicle and clozapine-n-oxide (CNO) injection. **(B)** Example of a heatmap of the time spent in each area of the SIM arena with a receptive female stimulus or male control stimulus rat on each side. **(C)** Example images of DREADD expression. Dotted lines delineate BNST. LV; lateral ventricle, aca; anterior commissure, 3V; third ventricle. **(D)** Number of c-Fos+ cells after homecage control condition (HC) or after copulation to one ejaculation (COP). (n=6) **(E)** Time spent in each of the stimulus zones in the SIM test. +p <0.05 compared to female zone. **(F)** Preference score (time spent in female zone/total time spent in female + male zones). +p < 0.05 compared to 0.50. **(G)** Time to performance of first copulatory behavior (i.e., mount or intromission). **(E, F, G)** n = 12 for Sham and Gi bars, n = 11 for Gq bars. **(All panels)** *p < 0.05

### Confirmation of DREADD expression and receptor activation by CNO treatment

Extensive expression of DREADD was observed throughout the anterior, lateral, ventral, and posterior BNST (Fig. 1C). Therefore, no animals were excluded from analysis based on DREADD expression. To assess whether CNO evoked increased BNST neuronal activity in the Gq-animals and decreased behavior-induced BNST neuronal activity in Gi-animals, we stained and quantified BNST cells positive for the neuronal activity marker c-Fos. We confirmed the effectivity of CNO treatment and DREADDs, as CNO significantly increased c-Fos+ cell density in all BNST subregions of Gq-animals compared to Sham animals (Suppl. Fig. 1A;), and decreased copulation induced c-Fos+ cell density in the posterior BNST of Gi-animals compared to Sham animals (Suppl. Fig. 1B).

### BNST neurons activated during copulation

First, we wanted to confirm the involvement of the BNST in sexual behavior. Therefore, we assessed copulation-induced c-Fos+ cell density in each of the subregions of the BNST (Suppl. Fig. 6-7). We found that copulation until one ejaculation induced an increase in c-Fos+ cell density in all BNST subregions in CNO-treated Sham animals as compared to homecage residing CNO-treated Sham animals (Fig. 1D).

### A modulatory role for BNST^CaMKIIa^ neurons in sexual incentive motivation

Next, we aimed to study the effects of silencing and stimulating the BNST on sexual incentive motivation. While the SIM test is designed to study sexual incentive motivation in particular, incentive motivation can also be interpreted from the latency to start copulation. Pair housed Sham, Gi-and Gq-animals were first placed in a 10-min SIM test with a control male and a receptive stimulus female as incentive stimuli, immediately followed by a 30-minute copulation test with a receptive stimulus female. We found that all subject males were sexually motivated, as they spent significantly longer in the female zone than in the male zone after VEH treatment (Fig. 1B, 1E). In addition, their preference scores were greater than 0.5 after VEH treatment (Fig. 1F).

Silencing the CaMKIIa+ neurons did not negate the presence of sexual motivation (Fig. 1E, 1F). However, it was found that silencing BNST^CaMKIIa^ neurons did change the magnitude of sexual motivation. Gi-animals spent less time in the female zone after CNO treatment compared to VEH treatment (Fig. 1E), and although following a similar pattern, this effect was not large enough to cause a significant effect on preference score (Fig. 1F). In addition, CNO also delayed the first copulatory behavior in Gi-animals compared to VEH in the copulation test (Fig. 1G) and compared to Sham-CNO. Altogether, this suggests that silencing CAMKII+ neurons in the BNST reduces the degree of sexual incentive motivation.

Interestingly, stimulating BNST^CaMKIIa^ neurons did not have the opposite effect of silencing. Just as with silencing, sexual incentive motivation was not negated by CNO treatment in Gq-animals (Fig. 1E, 1F), the magnitude of the motivation was not changed. If any effect, CNO actually reduced the preference score compared to VEH. However, the Sham group also spent more time in the female zone after receiving CNO compared to VEH. As CNO treatment did not change any other parameter in the sham-animals, nor did it on Gq-animals, these findings suggest that these altered outcomes on incentive motivation were probably artifacts.

Collectively, these findings indicate that while neural activity of CaMKIIa-neurons in the BNST is not essential for initiating motivation to engage in copulation, these neurons may exert a regulatory influence on sexual incentive motivation. Although silencing these neurons reduces the degree of sexual motivation, stimulating them does not increase the level of motivation in rats.

### Silencing BNST^CaMKIIa^ neurons affects sexual behavior, by inhibiting motivation to continue copulation

Next, we investigated the role of BNST^CaMKIIa^ neurons on copulation. Immediately when introduced to a receptive female, male rats perform several mounts and intromissions that proceed in rapid succession, and finally trigger an ejaculation. After a postejaculatory interval of male inactivity, the male rat starts a new ejaculation series cycle with copulatory behaviors (Fig. 2A, reviewed in (Heijkoop et al. 2018)). Recently, we used a new scoring scheme for evaluating male rat sexual behavior in more detail (Huijgens et al. 2021a). This advanced behavioral assessment of sexual behavior additionally allows for an in-depth analysis of the temporal patterning of mounts and intromissions in *mount bouts* and the breaks between them (*time-outs*). This temporal patterning, the intensity of mounts and intromissions, as well as the inter-copulatory interval durations, determine the efficiency of the copulatory behavior.

**Figure 2.**
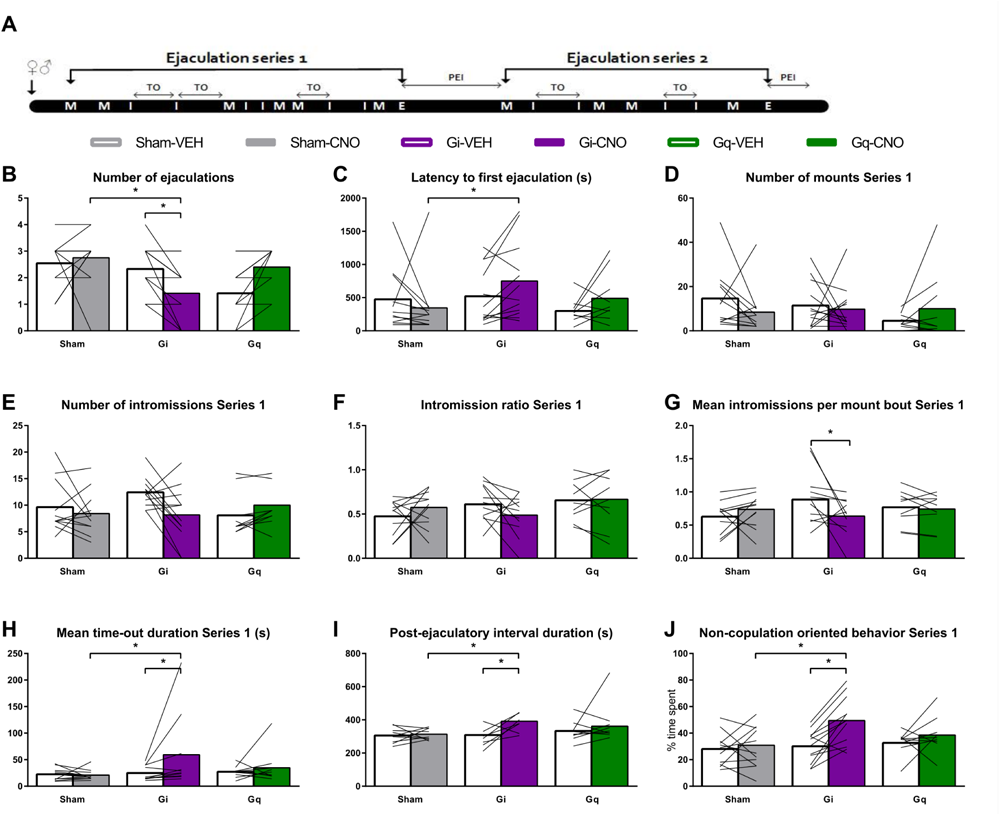
Effects of DREADD on copulation parameters. (A) Schematic representation of a sexual behavioral cycle of male rats with mounts (M), intromissions (I), ejaculations (E), and post-ejaculatory interval (PEI). The behaviors are divided into mount bouts interrupted by the time-outs (TO). (B) Total number of ejaculations in 30-minute test. (C) Latency to first ejaculation. (D) Number of mounts preceding the first ejaculation (Series 1). (E) Number of intromissions preceding the first ejaculation. (F) Intromission ratio (number of intromissions/number of mounts + intromissions) preceding the first ejaculation. (G) Mean number of intromissions in a mount bout in the first ejaculation series. (H) Mean duration of time-outs in the first ejaculation series. (I) Duration of the first post-ejaculatory interval (time from ejaculation to first copulatory behavior). (J) Percentage of time until first ejaculation spent on non-copulation-oriented behaviors (i.e., every behavior except for copulations, genital grooming, anogenital sniffing, chasing, and head directed to the female). (All panels) Left to right: n = 11, 12, 12, 12, 10, 10; (exception: (I) n = 10, 11, 11, 8, 10, 10). *p < 0.05

We found that inhibiting CaMKIIa+ neurons in the BNST significantly reduced the number of ejaculations (Fig. 2B) compared to both VEH and sham-CNO in a 30-minute copulation test. This decrease in number of ejaculations seems to be caused by an increase in ejaculation latency (Fig. 2C), rather than a reduced sensitivity to copulatory stimulations. In fact, silencing BNST^CaMKIIa^ neurons did not change the number of mounts (Fig. 2D) or intromissions (Fig. 2E), nor did the intromission ratio change (Fig. 2F; measure for copulatory efficiency, reviewed in (Heijkoop et al. 2018)). Notably, earlier studies have demonstrated an ejaculation deficit in BNST-lesioned male rats, accompanied by an increase in the number of intromissions preceding ejaculation. However, these lesion studies are likely confounded by additional lesions of the stria terminalis (Emery and Sachs 1976; Valcourt and Sachs 1979). When a detailed analysis of the temporal patterning was performed, we found that CNO treatment also did not change the number of mount bouts (Suppl. Fig. 2A), or mean number of mounts per mount bout (Suppl. Fig. 2B) in Gi-rats. Silencing the BNST^CaMKIIa^ neurons did, however, reduce the mean number of intromissions per mount bout (Fig. 2G), although this effect was not different from CNO-sham. Stimulating the BNST^CaMKIIa^ neurons did not affect any of the parameters mentioned above (Fig. 2 and Suppl. Fig 2).

The lack of effect on copulatory behaviors other than ejaculation suggests that CaMKIIa+ neurons in the BNST do not regulate copulatory sensitivity through sensory processing. Therefore, the increase in ejaculatory threshold must be caused by another regulatory mechanism. This suggestion was confirmed by our detailed behavioral analysis of the inter-copulatory intervals. Once copulation had commenced, chemogenetic silencing of BNST^CaMKIIa^ increased the latency to ejaculation through a slowing of the copulatory pace, as reflected in longer time-outs (Fig. 2H) and post-ejaculatory interval (PEI) durations (Fig 2I). We have previously shown that time-out and PEI durations are highly correlated within rats, leading us to hypothesize that these copulatory pauses may be regulated by a shared neural mechanism (Huijgens et al. 2021b). Altogether, we propose that the BNST might be a key regulator of copulatory pause durations.

In addition, a pattern of decreased interest in copulation arises from the data: silencing BNST^CaMKIIa^ decreased the relative time spent on genital grooming (Suppl. Fig 2C), and chasing after the female (Suppl. Fig 2D), and increased the time spent on non-copulation oriented behavior (Fig. 2J), which includes grooming other regions than the genitals (Suppl. Fig 2G) and head oriented away from the female (Suppl. Fig 2H). Finally, the non-copulation, female-oriented behaviors anogenital sniffing and head oriented towards the female remained unaffected (Suppl. Fig 2E, 2F). These effects on copulation and other behaviors during the copulation test are not likely to be caused by locomotor effects, as there were no CNO-induced effects on locomotor parameters (distance moved and velocity) in the SIM test (Suppl. Fig. 1C, 1D).

In summary, our data suggest that CaMKIIa+ neurons in the BNST are involved in the regulation of both initial motivation to engage in copulation, as well as in the motivation to continue copulation. The BNST^CaMKIIa^ neurons play an important role in regulating the temporal patterning of copulation to achieve an ejaculation and in modulating the motivational aspects of re-engaging after copulatory pauses.

### Overlap of regulation of sexual behavior and sucrose self-administration on the behavioral and neurobiological level

An important aim of the study was to assess if distinct micropatterns of reward seeking behavior overlap among diverse natural rewards and to determine the degree to which underlying neurobiological mechanisms converge. Therefore, in a separate sucrose self-administration experiment, we also chemogenetically silenced and stimulated the BNST^CaMKIIa^ neurons (Fig. 3A). After brain surgery and 4 weeks of recovery, subject males were trained and tested in operant conditioning chambers that were equipped with two retractable levers and a sucrose pellet dispenser.

**Figure 3.**
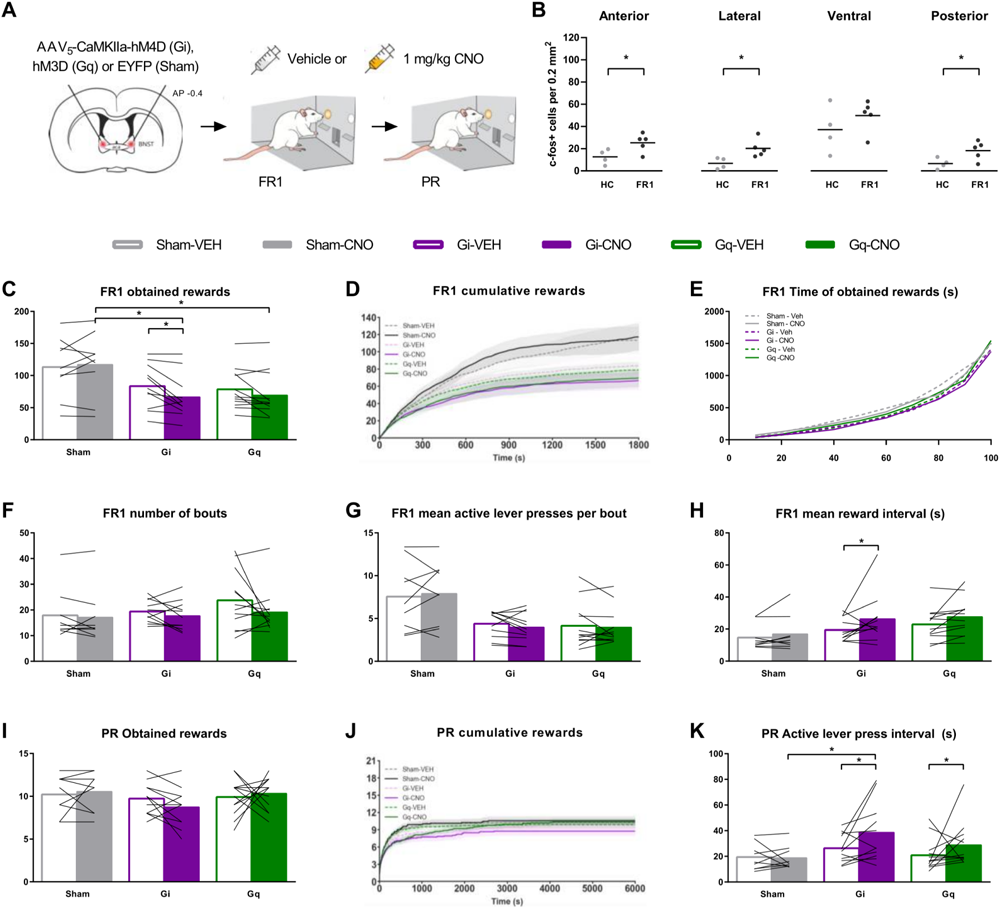
Effects of DREADD on sucrose self-administration. (A) Bilateral viral targeting of the bed nucleus of the stria terminalis (BNST) and experimental timeline. Subject males were tested in sucrose self-administration test under FR1 and PR schedules after vehicle and clozapine-n-oxide (CNO) injections. (B) Number of c-Fos+ cells after homecage control condition (HC) or after 30 minutes of FR1 engagement (FR1). (HC, n=4, FR1 n=5 Sham rats) (C) Number of obtained rewards during 30 min FR1 test. (D) Cumulative rewards in bins of 1 second, plotted as group mean ± s.e.m. in FR1 test. (E) Cumulative distribution of percentage of total obtained rewards over time in bins of 10% of total rewards in FR1 test. (F) Total number of lever pressing bouts in the FR1 test. (G) Average number of active lever presses per lever pressing bout in FR1 test. (H) Mean interval between rewards, calculated for the time between first and last reward in FR1 test. (I) Total number of obtained rewards under progressive ratio (PR). (J) Cumulative rewards in bins of 1 second, plotted as group mean ± s.e.m. in PR test (K) Mean interval time between active lever presses over the total duration of the PR test (C-K) n = 9 for Sham, 11 for Gi, 12 for Gq. Data points FR1 are averages of 2 vehicle-and 2 CNO-tests. (All panels) *p < 0.05

### BNST neurons are activated during sucrose self-administration

First, we confirmed the successful application of our methods and the involvement of the BNST in sucrose self-administration. The DREADD expression, location, and spread in and around the BNST in the sucrose self-administration experiment was similar to that of the animals in the sexual behavior experiment (Fig. 1C). No animals were excluded from the data analysis based on DREADD expression. Moreover, we again confirmed the effectivity of CNO treatment by treating Gq-animals with CNO in their homecage. We found that c-Fos+ cell density was increased in all subregions except for ventral BNST as compared to CNO-treated Sham animals (Suppl. Fig. 1E). In order to investigate the involvement of the BNST in sucrose self-administration, we assessed sucrose intake-induced c-Fos+ cell density in each of the subregions of the BNST. We found increased c-Fos+ cell density in all subregions except for the ventral BNST in CNO-treated Sham animals after 30 min of sucrose self-administration under FR1 as compared to CNO-treated Sham animals residing in the homecage (Fig. 3B). DREADD silencing of the BNST in the Gi group did not reduce the increase in c-Fos+ cells upon sucrose self-administration in any of the BNST subregions as compared to CNO-treated Sham animals (Suppl Fig.1F), which is probably a result of the already low c-Fos+ count in the control group.

### BNST^CaMKIIa^ neurons play a similar role in sucrose self-administration as in sexual behavior

To explore the effects of BNST^CaMKIIa^ neuronal modulation on reward sensitivity and motivation for sucrose self-administration, we first employed a fixed ratio 1 (FR1) paradigm. We found that silencing of the BNST reduced the number of rewards obtained under FR1 through a similar behavioral mechanism as during copulation: the pace of obtaining rewards was slowed down. CNO decreased the number of obtained rewards in Gi-rats compared to VEH (Fig. 3C) and compared to Sham-CNO, while the number of inactive lever presses remained unaffected (Suppl. Fig. 3A). Just as with sexual behavior, we found that as soon as the first rewards were obtained, silencing BNST^CaMKIIa^ neurons slowed down the rate of obtaining more sucrose rewards (Fig. 3D). Distributions of time to each next 10% of total obtained rewards showed that the overall pattern of obtaining rewards throughout the test was consistent (Fig. 3E), i.e., the rate of obtaining rewards was consistently faster in the beginning of each test and then slowed down for Gi-animals treated with both VEH and CNO (see also Suppl. Fig. 4). This suggests that inhibition of BNST^CaMKIIa^ neural activity does not necessarily affect incentive motivation, but decreases the tendency to continue to obtain more sucrose.

In sexual behavior, we found that the reduced number of ejaculations obtained was a result of longer pauses rather than reduced copulatory sensitivity. Although the conditioning chamber is less suitable for such detailed behavioral analyses, we attempted to make it as comparable as possible by identifying FR1 lever pressing bouts. Assuming that a distribution of all intra-bout interval durations has a smaller range and lower variance than inter-bout-intervals within animals, we ordered and plotted all lever press interval durations under vehicle conditions as scree plots for each animal (see suppl. Fig. 5 for example). Every scree plot was then visually examined to identify the approximate inflection point of the plot. This inflection point (10 seconds) was considered the threshold duration for inter-bout-intervals, used for the lever pressing bout analyses. Similar as with sexual behavior, there were no effects on the number of bouts (Fig. 3F), and average number of lever presses per bout (Fig. 3G). Interestingly, we found that the reduced number of rewards were indeed aligned with longer intervals between rewards, although this effect was not seen as longer pauses between bouts. CNO treatment increased the average duration of inter-reward intervals in Gi-animals compared to VEH (Fig. 3H).

By increasing the efforts for a reward, and thereby studying the motivation to continue obtaining rewards in more detail, rats were subsequently trained under a progressive ratio (PR) schedule of reinforcement (Richardson and Roberts 1996). We found that BNST^CaMKIIa^ neuron silencing did not alter the total number of rewards obtained (Fig. 3I), or active lever presses (Suppl. Fig. 3B) under a PR paradigm, nor did it change the breakpoint (Suppl. Fig. 3C) or number of inactive lever presses (Suppl. Fig. 3D). However, we did again observe a reduced pace of active lever pressing upon silencing of the BNST (Fig. 3J). This reduced pace was again reflected by an increased interval between active lever presses (Fig. 3K) without changing the total test duration (Suppl. Fig. 3E). Together, this confirms that inhibition of BNST^CaMKIIa^ neurons leads to lower levels of motivation to continue self-administering sucrose rewards, just as it reduced motivation for copulation. While it reduced the total number of rewards obtained under a FR schedule, it did not result in fewer rewards obtained under a PR regime. This may be explained by the fact that while FR1 sessions have a fixed duration, under a PR schedule of reinforcement, the rats have unlimited time to obtain rewards as long as they reach their threshold for the next reward within the set 30 minutes. Therefore, a reduced pace is more likely to affect the number of rewards under a FR1 schedule when compared to a PR schedule. However, since 90% of the rewards were collected within 15 minutes under the FR1 schedule, this seems an unlikely explanation for the present findings. Instead, we could speculate that the reduced engagement to continue acquiring rewards is more strongly affected by the BNST under a low-effort regime, and more subtly under a high-effort regime. This hypothesis is also supported by the lack of effect on sucrose intake found under a FR4 regime upon BNST silencing (Flanigan et al. 2023a) In any case, since the pace to obtain rewards is still affected under PR, these findings suggest that BNST^CaMKIIa^ neurons play a role in motivation under both low-and high-effort paradigms.

Just as with sexual behavior, stimulating BNST^CaMKIIa^ neurons did not really seem to change the motivation for sucrose. No changes were found in the total number of obtained rewards under both FR (Fig. 3C) and PR (Fig. 3I), or in the breaking point (Suppl. Fig. 3C) between VEH and CNO treatment in Gq-rats. There was a reduction in rewards obtained under FR1 compared to Sham-CNO (Fig. 3C), but this effect was not different from Gq-VEH. We did find an increase in active lever press intervals of Gq-CNO rats compared to Gq-VEH rats (Fig. 3K), but this effect was not different from Sham-CNO. Overall, we conclude that BNST^CaMKIIa^ neurons play a similar regulatory role in different natural rewarding behaviors: sexual behavior and sucrose self-administration. While silencing results in an inhibitory effect on both initial motivation to engage in copulation, as well as on the motivation to continue copulation, stimulation does not affect these parameters.

### Internal state of need modulates BNST^CaMKIIa^ regulation of sexual behavior

Finally, given the assumed role of the BNST in valence surveillance, weighing internal state and environmental stimuli, we tested whether manipulation of internal state (through social isolation or food restriction, respectively) affected behavioral outcomes of silencing and stimulating BNST^CaMKIIa^ on both sexual behavior and sucrose self-administration. If BNST^CaMKIIa^ is key to sensing the internal state in order to guide an appropriate response to a stimulus, we expected to see a blunted effect of deprivation during chemogenetic inhibition. In a sexual context, inducing a state of need would mean inducing a state of need for copulation, which we are not able to do experimentally. Therefore, we induced a state of social need through single housing. One week of single housing already greatly increases contact time and social interactions in a social interaction test (Latane et al. 1972; Niesink and Van Ree 1982) and two weeks of single housing has previously been shown to reduce the number of mounts, intromissions, and ejaculations and increase the latency to the first of these behaviors in a copulation test (deCatanzaro and Gorzalka 1979). For this study, we chose to single house the male rats for 1 week before each testing day (vehicle and CNO).

First, we did not find any effects of single housing in and of itself on sexual incentive motivation (Fig. 4A-C compared to Fig. 1E-G), nor on copulation (Fig. 4D-I compared to Fig. 2B,C, F-J). All rats showed normal levels of sexual motivation and behavior suggesting that even though 1 week of single housing affects social behavior, it might be too short to induce copulatory deficits or sexual experience could have annulled such effects. However, single housing did modulate the sexual effects of chemogenetic silencing BNST^CaMKIIa^. After single housing, sexual incentive motivation was now reduced upon BNST^CaMKIIa^ silencing in all parameters: both time spent in the female zone (Fig. 4A) and preference score (Fig. 4B) were decreased. Moreover, the increase in latency to 1^st^ copulatory behavior was persistent after single housing (Fig. 4C). Single housing by itself did not alter the magnitude of motivation upon BNST^CaMKIIa^ stimulation, except that the potential artifact of a reduced preference score during pair housing was removed. Altogether, we can conclude that social deprivation strengthens the effects of chemogenetic silencing of BNST^CaMKIIa^ on sexual incentive motivation.

**Figure 4.**
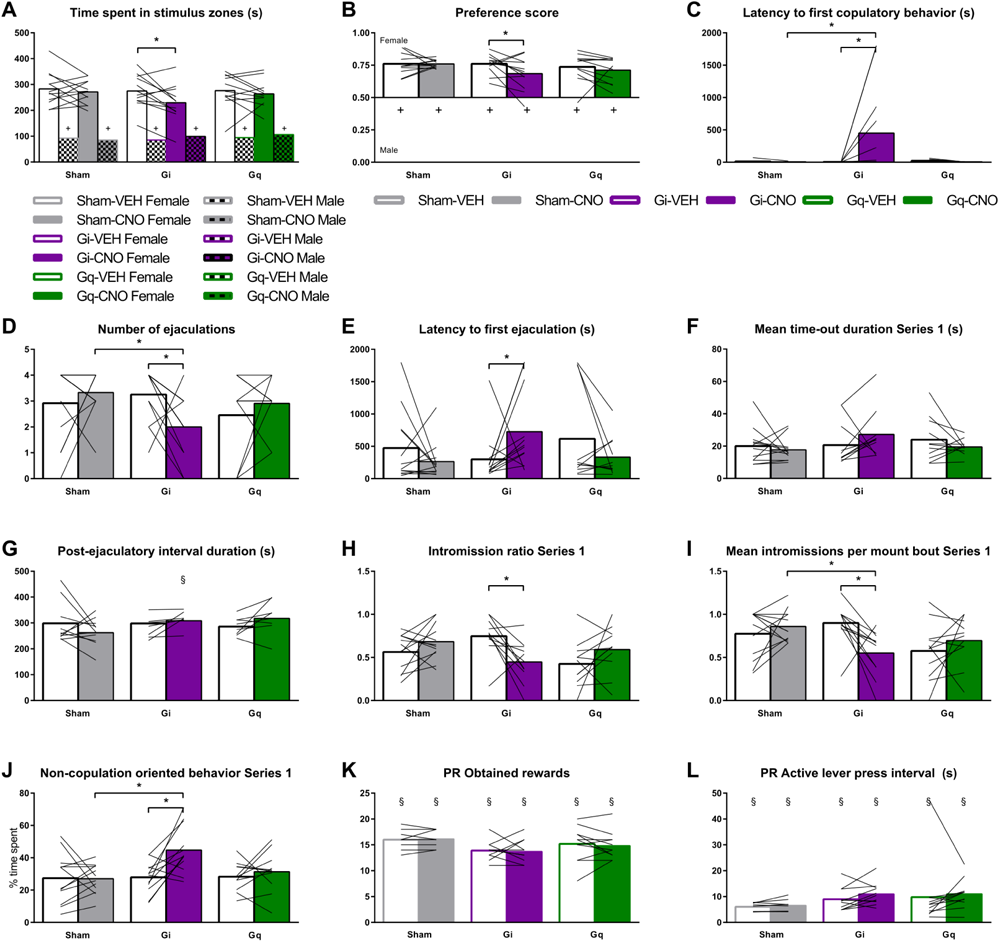
Effects of social-and food-deprivation on DREADD effect on sexual behavior and sucrose self-administration, respectively (A) Time spent in each of the stimulus zones in the SIM test. +p <0.05 compared to female zone. (B) Preference score (time spent in female zone/total time spent in female + male zones). +p < 0.05 compared to 0.50. (C) Time to performance of first copulatory behavior (i.e., mount or intromission). (D) Total number of ejaculations in 30-minute test. (E) Latency to first ejaculation. (F) Mean duration of time-outs in the first ejaculation series. (G) Duration of the first post-ejaculatory interval (time from ejaculation to first copulatory behavior). §p < 0.05 compared to pair housing condition (Fig.2I) (H) Intromission ratio (number of intromissions/number of mounts + intromissions) preceding the first ejaculation. (I) Mean number of intromissions in a mount bout in the first ejaculation series. (J) Percentage of time until first ejaculation spent on non-copulation oriented behaviors (i.e., every behavior except for copulations, genital grooming, anogenital sniffing, chasing, and head directed to the female). (K) Total number of obtained rewards under progressive ratio. (L) Mean interval time between active lever presses over the total duration of the PR test. (A-H.

During copulation, while no effects of BNST^CaMKIIa^ stimulation were found under single housing conditions, most BNST^CaMKIIa^ silencing-induced deficits that were found under pair housed conditions persisted after single housing. We still found that BNST^CaMKIIa^ silencing induced a reduction in number of ejaculations (Fig. 4D) as a result of longer latencies to ejaculation (Fig. 4E). Interestingly, though, while the BNST was involved in regulating copulatory behavior by controlling the copulatory breaks, social deprivation seems to attenuate this regulatory mechanism. We found that after single housing the increased time-out (Fig. 2H) and PEI (Fig. 2I) durations caused by BNST^CaMKIIa^ silencing in the non-deprived context, were no longer present (Fig. 4F and 4G respectively). In fact, the PEI of CNO-treated Gi animals under pair housing was longer than under single housing (Fig. 4G). Instead, effects on copulatory efficiency, i.e. decreased intromission ratio (Fig 4H), and mean intromissions per mount bout (Fig. 4I), are more pronounced after single-housing than after pair-housing. Thus, as the BNST^CaMKIIa^ silencing induced a deficit in initial sexual motivation prior to copulation persisted or even strengthened after single housing, deficits in the motivation to continue copulation may have been mitigated by social deprivation. However, time spent on non-copulation oriented behavior remained increased (Fig. 4J), just as the effects on other behaviors remained mostly unaffected by social deprivation (Suppl. Fig 2). This shows that animals did not increase their engagement level in copulation after single housing, nor did the rescued copulatory pace affect the ejaculatory deficits.

Overall, this suggests that BNST^CaMKIIa^ neurons are involved in the regulation of sexual behavior, regardless of the internal state of the animal. Still, an extraneously induced ‘internal state of need’, social deprivation, changes the role of these cells in motivated behaviors. While CaMKIIa-neurons in the BNST start to play a more pronounced role in modulating sexual incentive motivation when rats are socially deprived, they switch from controlling copulatory pauses to modulating copulatory efficiency. The role of BNST^CaMKIIa^ neurons in sexual behavior is thus – at least to some extent - influenced by the internal state of the animal.

### Internal state of need modulates BNST^CaMKIIa^ regulation of natural reward seeking behavior

To study the effects of the internal state on another natural reward seeking behavior, we tested the effects of food restriction on sucrose self-administration. Hereto, we removed the homecage food for 24 hours before re-testing under PR. Animals were tested twice after food restriction; once after CNO and once after VEH treatment (counterbalanced).

While the level of sexual motivation and behavior remained similar after social deprivation, food deprivation did alter sucrose self-administration parameters on a PR schedule. As expected, food-restricted rats increased the total number of rewards obtained (Fig. 4K compared to Fig. 3I)) and number of active lever presses (Suppl. Fig. 3B). As expected, food deprivation also decreased the average interval duration between active lever presses compared to *ad libitum* food access (Fig. 4L compared to Fig. 3K)), suggesting that food deprivation robustly increases the motivation for sucrose.

Interestingly, an internal state of hunger also attenuated the effects caused by BNST^CaMKIIa^ silencing or stimulating on the active lever press interval (Fig. 4L). Only the increased test duration in Gq-animals persisted after food restriction, but it was no longer different from sham animals (Suppl. Fig 3D). Finally, food restriction itself did not affect test duration and rats merely acquired more rewards in the same amount of time. Overall, this shows that the role of BNST^CaMKIIa^ neurons in sucrose self-administration depends on the internal state of the animal. While BNST^CaMKIIa^ neurons play a regulatory role in motivation via controlling the pace of reward taking under normal food conditions, these findings show that their impact becomes less pronounced after food deprivation. It could be argued that the increased levels of motivation and the intense willingness to work for sucrose rewards, induced by an internal state of hunger, reduces the active lever press interval to heights in which fluctuations caused by BNST^CaMKIIa^ silencing can no longer be observed. However, it could also simply be that the modulatory role of BNST^CaMKIIa^ neurons is replaced by a more pronounced role for other neurons in the BNST or elsewhere.

## Discussion

Overall, we propose that BNST^CaMKIIa^ neurons are involved in the temporal patterning of reward seeking behavior, primarily during “normal” physiological states, and during behaviors with a low effort-to-reward-ratio. Both FR1 and initial copulatory behavior require low effort from the subject animal to acquire the respective reward. In FR1, a reward equals a single lever press, and acquiring a single intromission in a copulation test only requires a short chase after the female in a small enclosure. The motivation to continue acquiring rewards is reflected in the duration of reward intervals in FR1, and copulatory pauses during copulation. This motivation to continue is affected by BNST^CaMKIIa^ silencing in all effort regimes. We could say that continuous copulatory behavior requires more effort than FR1, and the effort-to-reward-ratio is highest at late stages under PR. When more effort is required, BNST^CaMKIIa^ silencing results in longer active lever pressing intervals for sucrose rewards, which is comparable to the longer breaks seen in the copulatory test in both time-out and post-ejaculatory interval. This suggests that the role of BNST^CaMKIIa^ neurons converge on different types of natural rewarding behaviors.

With its tight connections with the medial amygdala (MeA)(Lebow and Chen 2016), it was expected that CaMKIIa-neurons in the BNST have a similar role in sexual behavior as CaMKIIa-neurons in the MeA. Previously, we have shown that silencing or stimulating MeA^CaMKIIa^ neurons does not affect sexual incentive motivation in rats (Huijgens et al. 2021c). Therefore, this study shows that CaMKIIa-neurons in the BNST and MeA play dissimilar roles in sexual motivation. Our data implies that the medial amygdala might regulate copulatory sensitivity and ejaculatory threshold through sensory processing (Huijgens et al. 2021d), while the BNST is involved in the temporal patterning of copulation, and the motivational aspects of re-engaging after copulatory pauses. Both affect the latency to ejaculation, but through different behavioral mechanisms.

Interestingly, no effects were found during BNST^CaMKIIa^ stimulation, congruent with previous results of MeA^CaMKIIa^ stimulation. In mice, chemogenetic stimulation of Esr2+ neurons in the BNST strongly suppresses sexual behavior (Zhou et al. 2023), whereas chemogenetic stimulation of aromatase+ neurons in the BNST promotes copulation (Bayless et al. 2019b). Moreover, Esr2+ optogenetic stimulation time-locked to anogenital sniffing suppresses the transition to mounting and reduces time spent on anogenital sniffing, whereas stimulation during intromissions does not interrupt normal copulation patterns. Since it is currently unknown whether CaMKIIa+ neurons in the rat BNST express Esr2 and/or aromatase, and what the complete transcriptome looks like, we cannot speculate much on the reasons for not finding an effect of CaMKIIa+ BNST neuron stimulation.

However, it is possible that continuously high levels of activity in these neurons is irrelevant, and that their effect is more dependent on more discrete bursts of activity in response to certain stimuli or behavior. Consequently, suppression of such bursting activity by chemogenetic silencing would then prevent these actions to take place.

Some limitations of this study should be addressed. First, there was extensive spread of DREADD expression to multiple brain regions adjacent to the BNST. In some animals, low density DREADD expression was also observed in a small medial portion of the caudate putamen, globus pallidus, ventral pallidum, and other surrounding regions. Future studies should confirm that all effects attribute to manipulation of the BNST, as some other regions expressing DREADD have been shown to be involved in socio-sexual behavior (Hull 2011), ingestive behavior (Rossi 2023; Stamos et al. 2023), and/or reward and motivation (Gordon-Fennell et al. 2019; Root et al. 2015; Sharpe et al. 2017). However, most DREADD was expressed in the BNST. In addition, as the BNST is closely connected to many brain regions including several cortical regions, the hypothalamus, amygdala, periaqueductal grey, ventral tegmental area and nucleus accumbens (Dong and Swanson 2004; Lebow and Chen 2016; Stamatakis et al. 2014), it is reasonable to assume that the BNST is not acting alone, but closely collaborates with other brain areas to regulate behavioral responses to different natural rewards. As mentioned previously, the BNST modulates reward and aversion through its projection to the ventral tegmental area (Jennings et al. 2013b; Soden et al. 2022), while connections with the amygdala, medial prefrontal cortex, and hypothalamus are involved in processing mood and arousal states and monitoring of food and fluid intake (Davis et al. 2010; Dong et al. 2001; Dong and Swanson 2004; Hammack et al. 2009; Radley and Sawchenko 2011; Rodriguez-Romaguera et al. 2020). Future research should reveal to what extent these projecting brain areas are involved in sexual behavior and sucrose consumption, and to what extent the underlying neurobiological mechanisms converge or overlap.

Finally, the transcription profile and anatomical connections of CaMKIIa+ neurons in the BNST are currently unknown. Even though the CaMKIIa promoter is often used to specifically target glutamatergic neurons, CaMKIIa promoter activity was shown in e.g. GABAergic neurons in several brain regions (Benson et al. 1992; Jennings et al. 2013a). Therefore, future research should address the nature of BNST^CaMKIIa^ neurons in more detail.

In conclusion, we found that BNST^CaMKIIa^ neurons are involved in the regulation of pacing both sexual behavior and sucrose intake. This suggests that different natural rewards could recruit similar brain circuitry to regulate the display of motivated behaviors. In addition, our study confirmed that the internal physiological state of the animal affects how the BNST modulates these behaviors.

Overall, this study offers important perspectives on the overlap of neural processes that drive the pursuit of different kinds of natural rewards and illuminates the interrelatedness of reward-associated behaviors in male rats.

## Methods

### Rats

In the sexual behavior experiment (carried out in Tromsø), 36 subject male Wistar rats, 10 stimulus female Wistar rats, and 4 stimulus male Wistar rats were acquired from Janvier Labs (France), and were pair housed (unless otherwise indicated) in Macrolon IV® cages on a reversed 12 h light/dark cycle (lights on between 23:00 and 11:00) in a room with controlled temperature (21 ± 1 °C) and humidity (55 ± 10 %), with ad libitum access to standard rodent food pellets and tap water throughout the experiment. In the sucrose self-administration experiment (carried out in Utrecht), a total number of 32 adult male Wistar rats, acquired from Charles River (Sulzfeld, Germany), was used. They were pair housed until brain surgery, and individually housed for the rest of the experiment in Macrolon III sawdust bedded cages on a reversed 12 h light/dark cycle (lights on between 07:00 and 19:00) in a room with controlled temperature (21 ± 2 C° and humidity (50-70%).

This study only investigated male behavior, because males and females show different kinds of sexual behaviors making it difficult to compare. We hope that future studies will address the same questions in female rats.

### Viral constructs and drugs

For sexual behavior, conducted at UiT The Arctic University of Norway (NO), AAV5-CaMKIIa-hM4D-mCherry (Gi; inhibitory DREADD, University of North Carolina Vector Core, Chapel Hill, USA), AAV5-CaMKIIa-hM3D-mCherry (Gq; stimulatory DREADDs, University of North Carolina Vector Core, Chapel Hill, USA) and AAV5-CaMKIIa-EYFP (Sham; no DREADDs, University of North Carolina Vector Core, Chapel Hill, USA) were used in this experiment. For sucrose self-administration, conducted at University of Utrecht (NL), the same viral constructs were used as for sexual behavior, but now obtained from Addgene, USA (pAAV5-CaMKIIa-hM4D(Gi)-mCherry and pAAV5-CaMKIIa-hM3D(Gq)-mCherry were gifted from Bryan Roth (Addgene viral prep # 50477and # 50476; pAAV-CaMKIIa-mCherry was a gift from Karl Deisseroth (Addgene plasmid # 114469). Viral constructs were used as is or diluted in saline before use, with a final viral titer between 1.4 and 4.3 x 10^12^ vg/mL. Clozapine N-oxide (CNO) (BML-NS-105; Enzo Life Sciences, Farmingdale, USA) was dissolved in ddH2O at a concentration of 1 mg/mL and kept at −20° until use. For experiments, rats were injected intraperitoneally with 1 mL/kg of the 1 mg/mL CNO solution or vehicle (ddH2O). Silastic capsules (medical grade Silastic tubing, 0.0625 in. inner diameter, 0.125 in. outer diameter; Degania Silicone, Degania Bet, Israel) for females were 5 mm long and contained 10% 17β-estradiol (E8875, Sigma, St. Louis, USA) in cholesterol (C3292, Sigma, St. Louis, USA). The silastic tubing was closed off by inserting pieces of toothpick into both ends and sealed with medical grade adhesive silicone (A-100; Factor II Incorporated, Arizona, USA). Progesterone (P0130, Sigma, St. Louis, USA) was dissolved in peanut oil (Apotekproduksjon, Oslo, Norway) at a concentration of 5 mg/mL.

### Surgical procedures

#### Brain surgery

##### Sexual behavior experiment

Brain surgery consisted of subsequent bilateral infusions of the viral vector solution into the BNST. Subject male rats were anesthetized with isoflurane and placed in a stereotaxic apparatus (Stoelting Europe, Ireland). A 10 uL Hamilton syringe with a 30G blunt needle was mounted in a Hamilton injector (Stoelting) on the stereotaxic apparatus and inserted into each brain hemisphere sequentially at AP -0.4mm, ML ±3.80mm, and DV -6.60mm in reference to bregma, at a 20° angle with respect to the DV-axis, inserting the needle tip in the coronal plane from the lateral aspect toward the medial aspect of each hemisphere. Per hemisphere, 250 nL of the viral construct solution was injected into the BNST at an infusion rate of 150 nL/min. Following infusion, the needle was left in place for 10 min before withdrawal and closing of the skin with a continuous intradermal suture (Vicryl Rapide 40, Ethicon, Cincinnati, USA). After surgery, rats were immediately pair housed. Analgesic treatment consisted of 0.05 mg/kg buprenorphine and 2 mg/kg meloxicam pre-operative and post-operative after 24 and 48 hours.

##### Sucrose self-administration experiment

The same volume of the same viral constructs was infused bilaterally into the BNST at the same coordinates during this experiment. Anesthesia consisted of a mixture of 75 mg/kg ketamine hydrochloride (Narketan 10%; VETOQUINOL, France) and 0.25 mg/kg dexmedetomidine (Dexdomitor; Pfizer Animal Health B.V., the Netherlands) injected intraperitoneally. The pressure injection was conducted using a 30G blunt cannula needle mounted in a holder on the stereotaxic apparatus (Kopf Instruments), attached to a piece of tubing (Plastics One, Roanoke, USA) connected to a Hamilton syringe mounted in a minipump. After viral construct infusions, a double guide cannula (26G, length 8.2 mm, ML distance 1.6 mm; Plastics One, Roanoke, USA) was implanted at AP -5.05mm, ML ±0.80mm, DV -7.15mm, targeting the ventral tegmental area, for intracerebral infusion of CNO to target the BNST → VTA projection. The guide cannula was fixed to the skull using stainless steel screws and surgical cement (Simplex™ P bone cement with tobramycin, Stryker Nederland B.V., The Netherlands). Anesthesia was terminated through subcutaneous administration of 1.0 mg/kg atipamezole (Antisedan; Pfizer Animal Health B.V., the Netherlands). After post-experimental brain processing revealed that the VTA guide cannulas were not in the target area. Therefore, the intracranial CNO infusion data were not further analyzed and omitted from this manuscript. After surgery, rats were single housed in individually ventilated cages for 7 days, and subsequently single housed in regular cages for the remainder of the experiment. Analgesic treatment consisted of 0.05 mg/kg buprenorphine and 2 mg/kg meloxicam pre-operative and post-operative after 24 and 48 hours.

##### Ovariectomy

Stimulus females for the sexual behavior experiment were ovariectomized under isoflurane anesthesia as previously described (Ågmo 1997). Briefly, a medial dorsal incision of the skin of about 1 cm, and small incisions in the muscle layer on each side, were made. The ovaries were located and extirpated, and the muscle layer sutured. A silastic capsule containing β-estradiol was placed subcutaneously. The skin was closed with a wound clip. Females were allowed at least one week of recovery before use in behavioral tests. Analgesic treatment consisted of 0.05 mg/kg buprenorphine and 2 mg/kg meloxicam pre-operative and post-operative after 24 and 48 hours.

### Behavioral assessment

#### Sexual incentive motivation

The sexual incentive motivation (SIM) test (see also: (Huijgens et al. 2023)) apparatus is a rectangular arena (100 × 50 × 45 cm) with rounded corners. At each long side, in opposite corners, a closed incentive stimulus cage is attached to the arena and separated from the arena by wire mesh (25 × 25 cm). Testing always takes place in a dimly lit (ca 5 lx) room. During testing, a social stimulus (intact male rat) was placed in one of the stimulus cages and a sexual stimulus (receptive female rat; ovariectomized, implanted with a 10% 17β-estradiol silastic capsule, and injected with 1 mg progesterone 4 h before use in the test (standard procedure in our laboratory (Huijgens et al. 2021a; Huijgens et al. 2021c; Ågmo and Snoeren 2017)) was placed in the other stimulus cage. The position of the stimulus cages and the stimulus rats were randomly changed throughout each experimental session. To start the test, the subject male was introduced to the middle of the arena and then video-tracked by Ethovision software (Noldus, Wageningen, the Netherlands) for 10 min. Virtual stimulus zones in (30 x 21 cm) were defined in the arena in front of each stimulus cage in Ethovision. The subject male was considered to be in one of the zones whenever its point of gravity was. Frequency of entering, and time spent in each of the zones, distance moved, and mean velocity, were output measures of Ethovision. The preference score was calculated by dividing the time spent in the female incentive zone by the total time spent in incentive zones. Between tests, feces and urine were removed from the arena, and the SIM arena was cleaned with diluted acetic acid between experimental days.

#### 2.4.2. Copulation

The copulation test was conducted in rectangular boxes (40 × 60 × 40 cm) with a Plexiglas front, in a dimly lit (ca 5 lx) room. At the time of testing, the male subject was allowed 5 minutes of habituation to the copulation box, after which a receptive female (injected with 1 mg progesterone 4 h before use) was introduced, which started the 30 minute test. All copulation tests were recorded on camera and behavior was later assessed from video. Behavioral assessment consisted of scoring behavioral events in Observer XT software (Noldus, Wageningen, the Netherlands). For the entire 30 min, 100% of the elapsed time was behaviorally annotated, similar to previous experiments (Huijgens et al. 2021a; Huijgens et al. 2021c). The ethogram consisted of the copulatory behaviors mount (pelvic thrusting while being mounted on the female), intromission (mounting the female with pelvic thrusting and penile insertion into the vagina; characterized by a more vigorous dismount) and ejaculation (characterized by a longer intromission and the female moving away from the male), as well as genital grooming (autogrooming of the genital region), other grooming (autogrooming in other regions than the genitals), chasing (running after the female), anogenital sniffing (sniffing the anogenital region of the female), head towards female (head oriented in the direction of the female while not engaging in other behavior, also includes sniffing the female), head not towards female (any behavior that is not oriented towards the female except grooming, e.g. walking, sniffing the floor, standing still with head direction away from female). From these data points, the following outcome measures were determined: number of ejaculations, number of mounts, intromissions, and mount bouts, intromission ratio (number of intromissions divided by number of mounts + intromissions), average mounts and intromissions per mount bout, mean time-out duration, post-ejaculatory interval duration (time from ejaculation to first mount or intromission), latency to ejaculation (time from first mount or intromission to ejaculation), latency to first behavior (time from start of the test to first mount or intromission), and percentage of time spent on behaviors. The outcome measures were calculated for the first ejaculation series (referred to as “Series 1”), except for the total number of ejaculations in the 30 min test. A mount bout is defined as “a sequence of copulatory behaviors (one or more), uninterrupted by any behavior (other than genital autogrooming) that is not oriented towards the female” (Huijgens et al. 2021b; Sachs and Barfield 1970). Mount bouts were identified by reviewing the events between each copulatory behavior (i.e. mount or intromission) using a python script (available upon request). Whenever “other grooming” or “head not towards female” occurred in between copulatory behaviors, this marked the end of the previous mount bout (end time was then set on the end of the last copulatory behavior) and the beginning of the next mount bout (start time of the next copulatory behavior), and the time in between these mount bouts as a time out. Time spent on non-copulation oriented behavior was defined as time spent on (non-genital) grooming + head not towards female.

#### Operant sucrose self-administration

##### Apparatus

Subject male rats were trained and tested in operant conditioning chambers (29.5 x 24 x 25 cm, Med Assiociates Inc., USA) in light-and sound-attenuating boxes with a metal grid floor (bars 1.57 cm apart), a white house light (28V, 100 mA) and a ventilation fan, controlled by Med-PC IV software (version 4.2). The chambers were equipped with two retractable levers (4.8 x 1.9 cm), a white cue light (28 V, 100 mA) above each lever, and a sucrose pellet receptacle (equipped with an infrared beam for nose poke detection) underneath a sucrose pellet dispenser in between the two levers. One lever was designated as “active”, responding on which was reinforced with a sucrose pellet (45 mg; TestDiet, USA), the other lever was designated as “inactive”, responding on which was recorded but initiated nothing. The position of the active and inactive levers was counterbalanced between rats and within groups.

##### Fixed ratio 1

Subject male rats were trained to respond for sucrose pellets during 30-minute operant sessions, once daily, 2-4 times a week. All rats had a minimum of 12 training sessions before testing. Some rats received 1 or 2 extra training sessions to achieve stable active lever pressing performance. Performance was considered stable at group level when the average number of active lever presses on individual days fell within a 90%-110% variability range of the total average of active lever presses over the last 3 training days. During training and testing, pressing the active lever turned on the cue light, dispensed a single sucrose pellet into the receptacle, retracted the levers, and turned off the house light. As soon as the animal nose-poked into the receptacle to retrieve the sucrose pellet, detected by interruption of the infrared beam, a new trial was started. Upon the start of a new trial, the levers were reintroduced, the cue light was turned off, and the house light was turned on again. The software output contained timestamps of each active lever response, inactive lever response, and first nosepoke after active lever response. This enabled analysis of response frequencies as well as latencies.

##### Progressive ratio

After FR1 training and testing, rats were trained under a progressive ratio (PR) schedule of reinforcement, where progressively additional active lever presses were required (response requirements: 1, 2, 4, 6, 9, 12, 15, 20, 25 etc. (Richardson and Roberts 1996)) for the subsequent single sucrose pellet reward. All subjects were trained once daily on 6 days over a period of 8 days. Performance was considered stable at group level when the average number of obtained rewards on individual days fell within a 90%-110% variability range of the total average of obtained rewards over the last 3 training days. A PR session ended when a rat failed to obtain a subsequent reward within 30 minutes.

### Brain processing, immunostaining and imaging

#### Brain processing

At the end of the experiments, rats were injected intraperitoneally with a lethal dose of pentobarbital (100 mg/kg), and transcardially perfused with 0.1 M phosphate buffered saline (PBS; pH 7.4) followed by 4% formaldehyde in 0.1 M PBS. Brains were removed and post-fixed in 4% formaldehyde for 48 hours. Subsequently, brains were kept in 20% sucrose in 0.1 M PBS until sunken, and then in 30% sucrose in 0.1 M PBS until sunken. Brains were then snap frozen in isopentane cooled by dry ice and kept in −80 °C until sectioning on either a cryostat (Cryostar NX70, Thermo Fisher Scientific, Waltham, USA), or a vibratome (Leica VT1200s) into 40 µm thick coronal sections and stored in cryoprotectant solution (30% sucrose w/v, 30% ethylene glycol v/v in 0.1 M PBS, pH 7.4) until further use.

#### Immunostaining

For immunohistochemistry staining of the DREADD-fused fluorophore, 1 in every 5^th^ free-floating brain section throughout and surrounding the BNST was washed in 0.1 M Tris-buffered-saline (TBS), blocked for 30 minutes in 0.5% BSA in TBS, and incubated on an orbital shaker for 24 hours at room temperature + 24 hours at 4 °C in polyclonal rabbit anti-mCherry (1:30 000, Abcam, cat. ab167453) or polyclonal chicken anti-EYFP (1:100 000), Abcam, cat. ab13970) antibody solution containing 0.1% Triton-X and 0.1% BSA in TBS. Sections were then incubated in biotinylated goat anti-rabbit (1:400, Abcam, cat. ab6720) or biotinylated goat anti-chicken (1:400, Abcam, cat. ab6876) antibody solution containing 0.1 % BSA in TBS for 30 min, avidin-biotin-peroxidase complex (VECTASTAIN ABC-HRP kit, Vector laboratories, cat. PK-6100, dilution: 1 drop A + 1 drop B in 10 mL TBS) solution for 30 min, and 3,3′-diaminobenzidine solution (DAB substrate kit (HRP), Vector laboratories, cat. SK-4100, dilution: 1 drop R1 (buffer solution) + 2 drops R2 (3,3′-diaminobenzidine solution) + 1 drop R3 (hydrogen peroxide solution) in 5 mL water) for 5-10 min, with TBS washes between all steps. Slides were dehydrated, cleared, and coverslipped using Entellan mounting medium (Sigma, St. Louis, USA). Immunohistochemical staining of c-Fos was carried out separately, on different brain sections than the DREADD fluorophore staining. Macroscopically matching AP anatomical location across brains, six sections (2 sections containing anterior and ventral BNST, 2 sections containing lateral BNST, and 2 sections containing posterior BNST) were selected for each animal. The same staining protocol as described above was employed, but a rabbit anti-c-Fos polyclonal antibody (1:4 000, Cell signaling, cat. 9F6) was used for primary antibody incubation.

#### Imaging

For imaging, slides were loaded into an Olympus VS120 virtual slide microscope system. High resolution image scans of the entire brain sections were obtained using a 10x objective using automatic focusing. For DREADD images, automatic exposure settings were used, whereas one and the same manual exposure time was used for c-Fos images across all animals and sections.

### Image analysis

#### DREADD

Using OlyVIA online database software (Olympus, Tokyo, Japan), the extent and location of DREADD expression was determined in each rat. DREADD expression in the BNST and in surrounding regions was categorized based on the amount of DREADD+ cells per region in the sample of sections (1 in 5 throughout the BNST). We qualified expression using a scoring system per brain region per hemisphere: 0 (no expression in the brain region); 1 (low expression; i.e. no more than a few positive cells in the region), 2 (medium expression throughout the brain region, typically >10 and <30 DREADD+ cells per section, or high expression in part of the brain region); and 3 (high expression throughout the brain region, typically >30 positive cells per section). A second observer validated the qualifications in 5 brains with various expression patterns. We then added the scores for each hemisphere and excluded rats with a total score (left + right hemisphere) smaller than 3 for the BNST from further analysis. Some rats in the Sham group (n = 6 in the total experiment) had very low DREADD expression and did not meet inclusion criteria. These animals were not excluded from data analysis, reasoning that lack of DREADD expression does not disqualify their control condition.

Furthermore, besides extensive expression of DREADD throughout the anterior, lateral, ventral, and posterior BNST (Fig. 1C), DREADD expression was also observed in brain regions ventro-posterior to the BNST in all animals, namely the reticular thalamic nucleus, the sublenticular extended amygdala, posterior lateral hypothalamus, and the lateral preoptic area. As the overall weight of expression was in the BNST, no animals were excluded from analysis based on DREADD expression.

#### c-Fos+ cell analysis

Images were loaded into FIJI ImageJ and 4 regions of interest were defined: anterior BNST (between AP 0.0mm and -0.12mm, dorsal of anterior commissure), ventral BNST (between AP 0.0mm and - 0.12mm, ventral of anterior commissure), lateral BNST (between AP -0.24mm and -0.36mm, lateral of anterior commissure), and posterior BNST (between AP -0.72mm and -0.9mm, between internal capsule and fornix). Boxes of always the same shape, size, and location were placed within these regions on both sides of the brain section images (one section per region of interest per brain, Suppl. Fig. 6). The image was converted to 8-bit and then to binary by thresholding to the same threshold for each image in same experiment. The threshold was determined by testing on a wide range of images with high and low c-Fos+ density. The ImageJ particle analyzer (size range 5-2000 pixels, circularity range 0.5-1.0) was used to count the number of particles (c-Fos+ cells) within each box on the binary thresholded image (Suppl. Fig. 7). For each animal, the number of c-Fos+ cells were averaged across hemispheres for each region of interest. The automated counts were validated by comparing to manual counting of c-Fos+ cells of a sample of boxes with a wide range of c-Fos+ density.

### Experimental design and interventions

#### Sexual behavior experiment

##### Pair housing

Subject males were first sexually trained in the copulation test set-up. Sexual training sessions consisted of allowing the males to copulate with a female until ejaculation, or for 30 minutes if no ejaculation was reached. When males had reached ejaculation in 2 sessions, a final full 30 minute copulation test (allowing for more than 1 ejaculation) was conducted for complete sexual experience. This ensured that each subject had ejaculated at least 3 times before experimental behavioral tests were conducted. Based on the number of ejaculations in the final training session, males were divided over 3 homologous phenotype groups for brain surgery. After brain surgery and recovery for 3 weeks, in the week before the first behavioral tests, subject males were habituated to the SIM arena without stimulus rats present in 3 sessions of 10 minutes. On subsequent testing days, subject males were first tested in the SIM test, 30 minutes after i.p. injection with 1 mg/kg CNO or vehicle, and then in the copulation test 5-15 minutes after the SIM test. Testing occurred once a week, following a Latin square within-subject design for balanced order of CNO and vehicle administration.

##### Single housing intervention

In the second part of the experiment, all males were single housed for 1 week before each testing day (vehicle and CNO) in order to study the effects of social deprivation.

For single housing, males were housed in smaller Macrolon cages with metal wire lids, still allowing for odors and sounds of rats in the same room to enter the cages. After single housing and the first test day, the subject males were rehoused in pairs for 1 week before being single housed for 1 week again for the second test day. Order of vehicle and CNO testing was counterbalanced within groups.

All behavioral tests were conducted during lights-off time.

#### Sucrose self-administration experiment

##### Ad libitum food access

After brain surgery and recovery, subject males were trained under FR1 until stable performance. Animals were subsequently tested 4 times under FR1; twice after CNO and twice after vehicle following a Latin square within-subject design. On testing days, rats were injected intraperitoneally with vehicle or CNO, and put in the operant box 30 minutes later. There was a minimum of 3 days between testing days, and each test was preceded by a baseline FR1 training session (no injections administered) the day before testing. After FR1 testing was completed, rats were trained under PR and tested 2 times; once after CNO and once after vehicle (counterbalanced). There was one day between the tests, and each test was preceded by a baseline PR training session the day before testing.

##### Food restriction

Rats underwent regular PR training sessions until re-testing under food restriction conditions. Food restriction consisted of removing homecage food access for 24 hours before testing under PR. Animals were tested twice after food restriction; once after CNO and once after vehicle (counterbalanced). There were 5 days between the tests, and each test was preceded by a baseline PR training session two days (to allow for the 24-hour food restriction) before testing.

All behavioral tests were conducted during lights-off time.

### Perfusion and DREADD validation

All rats were injected with 1 mg/kg CNO intraperitoneally on the day of euthanasia. Gq-animals, and half of the Sham-animals remained in the homecage after CNO injection and underwent cardiac perfusion with 4% formaldehyde 90 minutes later. In the sexual behavior experiment, all Gi-animals, and the other half of the Sham-animals were allowed to copulate until one ejaculation 30 minutes after CNO injection and were perfused 45 minutes after ejaculation. In the sucrose self-administration experiment, all Gi-animals, and the other half of the Sham-animals were allowed a 30-minute FR1 session in the operant box 30 minutes after CNO injection, were then returned to the homecage and perfused 30 minutes later. Brains of all animals were harvested for immunohistochemical analysis of DREADD expression and c-Fos protein positive cells.

### Data analysis and statistics

During copulation behavioral annotation from video, it was noticed that one of the females was not receptive on one of the test days. This female was paired with 3 subject males in the copulation test and no copulation occurred in these tests due to the lack of receptivity. The data from these 3 copulation tests have therefore been removed from the analysis. If a rat achieved no ejaculations in a 30-minute copulation test, the latency to ejaculation was scored as 1800 seconds (30 minutes).

Similarly, if a rat did not perform any copulatory behaviors, the latency to first behavior was scored as 1800 seconds.

Because the sucrose self-administration outcome measures under FR1 were quite variable over sessions within rats, each rat was tested twice after VEH and CNO treatment, and the data presented here are the comprised of the average of the two tests for each rat. Custom python scripts were used to extract the timestamps and latencies, and to calculate and create the cumulative curves of obtained rewards under FR1 and PR. In FR1, “mean reward interval” was calculated as (time of last reward – time of first reward)/ (number of obtained rewards -1). In PR, mean lever press interval was calculated as (total test duration)/(total number of active lever presses).

All data was analyzed in SPSS statistical software (IBM, version 29, Armonk, USA) and GraphPad Prism (version 9.5). A linear mixed model comprised of only the factor virus*treatment interaction term was run for each of the separate outcome measures of the FR1 sucrose self-administration under ad libitum food conditions. In case of a significant interaction effect (alpha 0.05), Bonferroni-corrected post hoc tests were conducted to identify significant within-(CNO vs. vehicle) and between-(only Gi vs. Sham and Gq vs. Sham) group differences. For the sucrose self-administration under PR and the sexual behavior outcome measures, a second interaction term factor of virus*treatment*condition (the latter being food restriction or single housing) was added to the linear mixed model. Appropriate Bonferroni corrected post hoc tests were run upon significant interaction (alpha 0.05). The SIM preference score was compared to chance (0.5) with a one-sample t-test for each condition within each treatment within each group. Time spent in female zone was compared to time spent in male zone, and female zone frequency was compared to male zone frequency, with a paired t-test.

In addition, we identified FR1 lever pressing bouts by assuming that a distribution of all intra-bout interval durations has a smaller range and lower variance than inter-bout-intervals within animals. Hence, we ordered and plotted all lever press interval durations under vehicle conditions as scree plots for each animal (see Suppl. Fig. 5 for example). Every scree plot was then visually examined to identify the approximate inflection point of the plot. This inflection point was considered the threshold duration for inter-bout-intervals. Since this threshold was around 10 seconds for most animals, we decided to use a 10 second threshold on all of the data for lever press bout analysis.

Densities of c-Fos+ cells were compared by means of 3 one-tailed independent samples t-tests (Gi vs. its Sham control, Gq vs. its Sham control, and the 2 Sham groups vs. each other, alpha=0.05), or one-tailed Mann-Whitney U test in case an F-test revealed unequal variances. For 2 animals in the sexual behavior experiment, data from the anterior and ventral BNST were excluded because of damaged sections prohibiting reliable counting of c-Fos+ cells.

## Acknowledgements

Financial support was received from the Research Council of Norway; grant #251320 to ES. We thank Carina Sørensen, Lorenzo Ragazzi, Ragnhild Osnes, Remi Osnes and Siri Knudsen for the excellent care of the animals in Tromsø, and Nicky van Kronenburg in Utrecht. In addition, we thank Anne-Marie Baars, José Lozeman-van ‘t Klooster, and Eline Elshof for their help with surgeries, and sucrose self-administration training and tests, and perfusions for the sucrose study. Finally, we thank the Advanced Microscopy Core Facility of UiT The Arctic University of Tromsø for access to their equipment for the brain analyses.

## Author contributions

PH: Experimental design, data gathering, behavioral annotation, methodology, data curation, analysis, writing -original draft. RH: Experimental design, methodology, writing – review and editing. HL: Experimental design, methodology, writing – review and editing. LV: Experimental design, methodology, writing – review and editing. ES: Experimental design, methodology, supervision, writing - original draft, funding acquisition.

