## Supplemental information for "CaMKIIa+ neurons in the bed nucleus of the stria terminalis modulate pace of natural reward seeking depending on internal state"

### Supplemental Figure 1

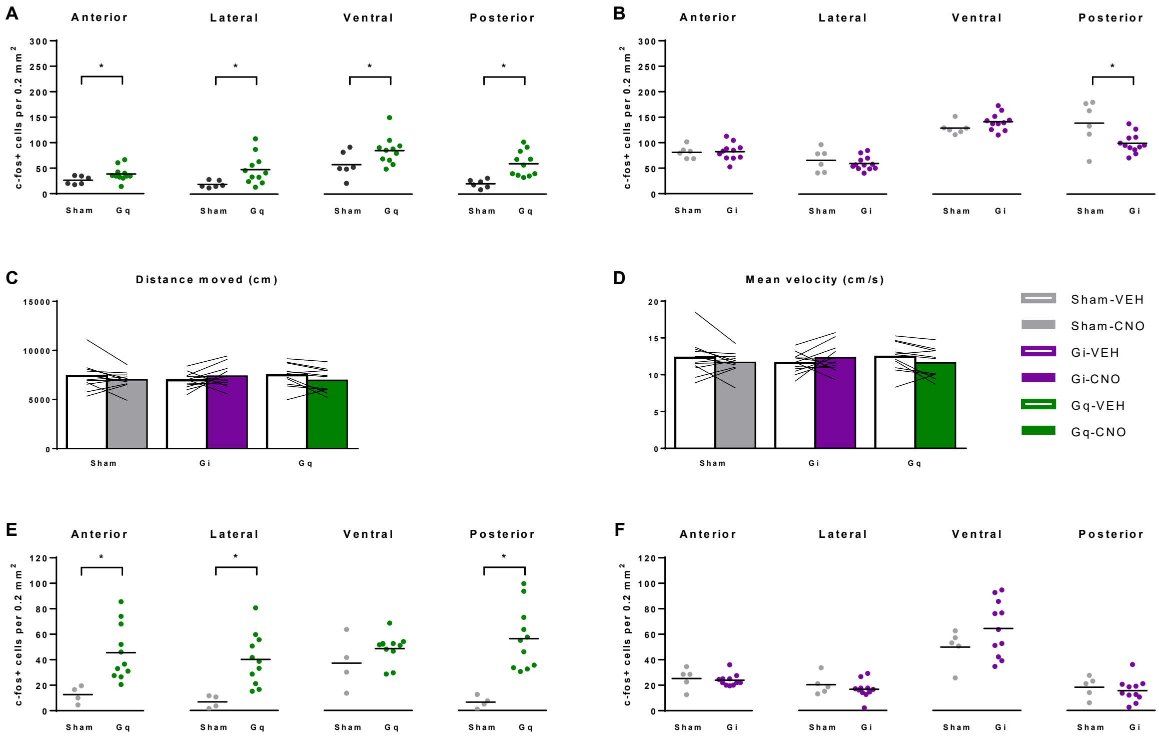


***Supplementary Figure 1. Effects of CNO treatment on c-Fos+ cell count and locomotor activity (A)*** *Number of c-Fos+ cells after CNO injection of Sham (n=6) or Gq-rats (n=11) in homecage of the sexual behavior experiment. (The Sham rats are the same rats as Fig. 1D-HC)* ***(B)*** *Number of c-Fos+ cells after CNO injection of Sham (n=6) or Gi-rats (n=11) after copulation to one ejaculation of the sexual behavior experiment. (The Sham rats are the same rats as Fig. 1D-HC)* ***(C)*** *Total distance moved during SIM test. n = 12 for Sham and Gi bars, n = 11 for Gq bars.* ***(D)*** *mean velocity during the SIM test. n = 12 for Sham and Gi bars, n = 11 for Gq bars.* ***(E)*** *Number of c-Fos+ cells CNO injection of Sham (n=4) or Gq-rats (n=11) in homecage of the sucrose experiment. (The Sham rats are the same rats as Fig. 3D-HC)* ***(F)*** *Number of c-Fos+ cells after CNO injection of Sham (n=5) or Gi-rats (n=11) after 30 minutes of FR1 engagement the sucrose experiment. (The Sham rats are the same rats as Fig. 3D-FR1)* ***(All panels)*** **p < 0.05.*

### Supplemental Figure 2

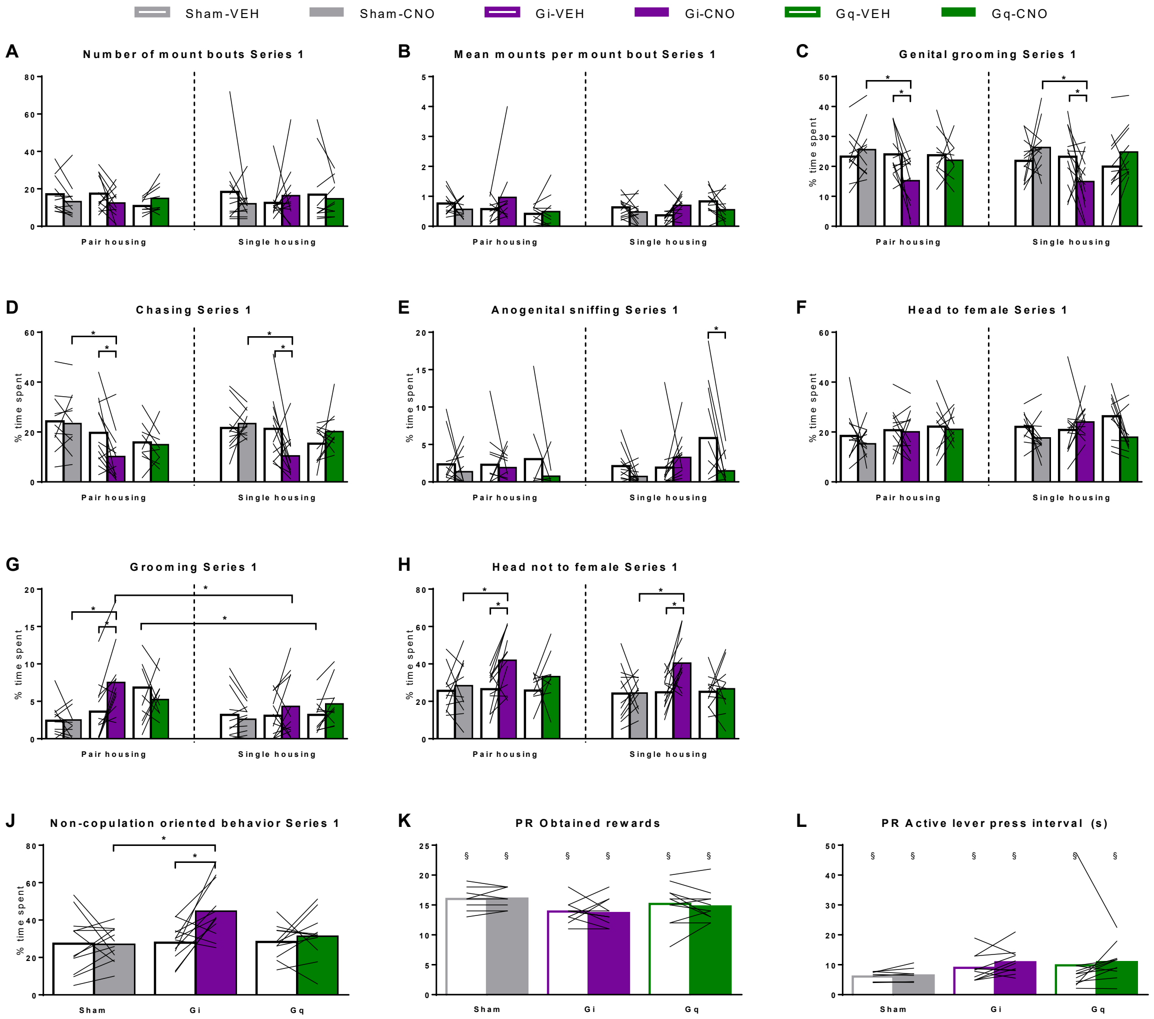


***Supplementary Figure 2. Effects of DREADD on copulation parameters of pair or single housed rats. (A)*** *Number of mount bouts preceding the first ejaculation (Series 1).* ***(B)*** *Mean number of mounts in a mount bout in the first ejaculation series.* ***(C)*** *Percentage of total time before the first ejaculation spent on genital grooming,* ***(D)*** *chasing,* ***(E)*** *anogenital sniffing,* ***(F)*** *having head in the direction of the female,* ***(G)*** *grooming of other regions than the genitals, and* ***(H)*** *Having head not in the direction of the female.* ***(All panels)*** *Left to right: pair housing n = 11, 12, 12, 12, 10, 10; single housing n = 12, 12, 12, 12, 11, 11. *p < 0.05. Statistical details of pair housing can be found in manuscript.*

### Supplemental Figure 3

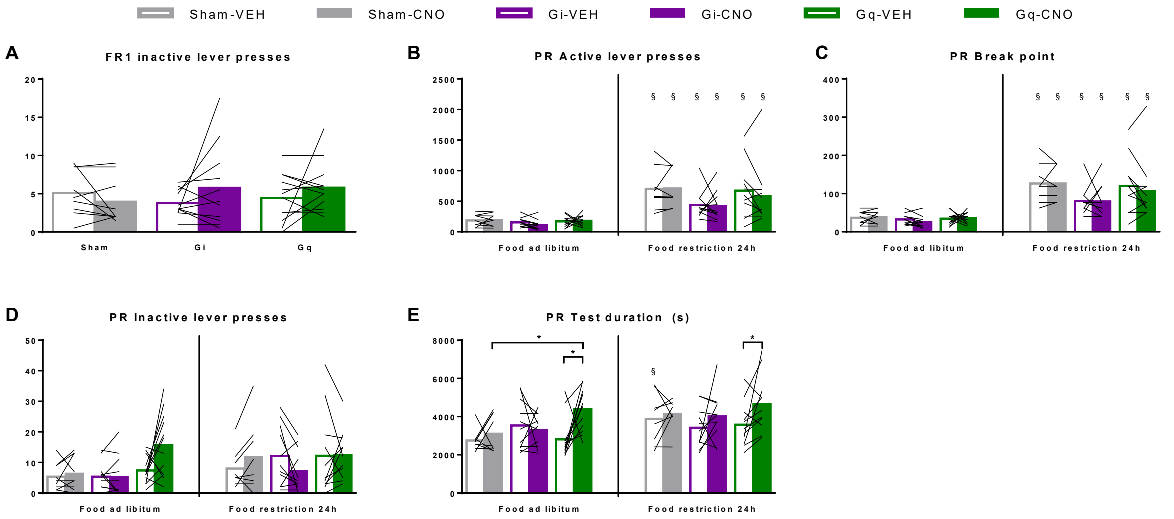


***Supplementary Figure 3. Effects of DREADD on FR1 and PR sucrose self-administration with food available ad libitum (FR1 & PR) or after 24 hours of food restriction (PR). (A)*** *Number of inactive lever presses under FR1 in the total 30 minute test. n = 9 for Sham, 11 for Gi, 12 for Gq. Data points are averages of 2 vehicle- and 2 CNO-tests.* ***(B)*** *Total number of active lever presses during the PR test. §p < 0.05 compared to food ad libitum condition.* ***(C)*** *Break point defined by the maximum number of lever presses made to earn the last reward.* ***(D)*** *Number of inactive lever presses under PR.* ***(E)*** *Total duration of the PR test (test ends after 30 minutes of no rewards obtained).* ***(B, C, D, E)*** *Ad libitum food n = 9 for Sham, 11 for Gi, 12 for Gq; Food restriction n = 8 for Sham, 11 for Gi, 12 for Gq. *p <0.05. Statistical details not mentioned here can be found in manuscript.*

### Supplemental Figure 4

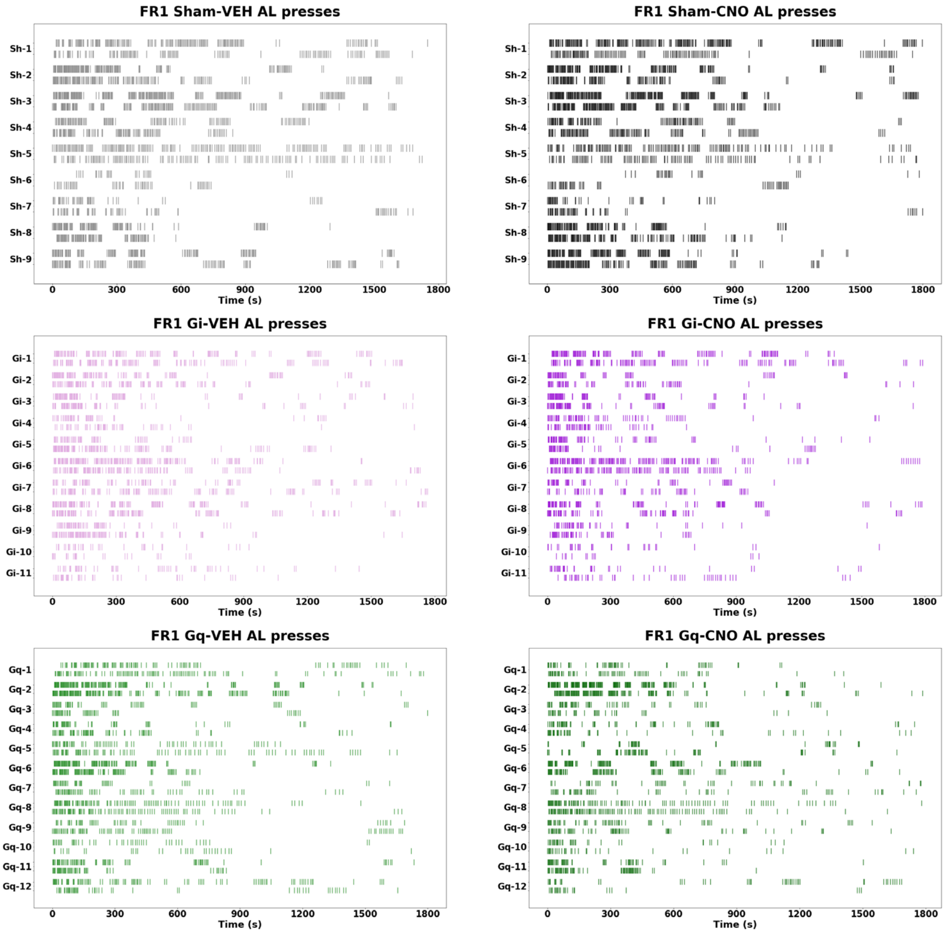


***Supplementary Figure 4.*** *Event plots of FR1 active lever presses. Every line represents an active lever press timestamp. Each animal was tested twice after vehicle and twice after CNO and these tests are plotted separately.*

#### Supplemental Figure 5

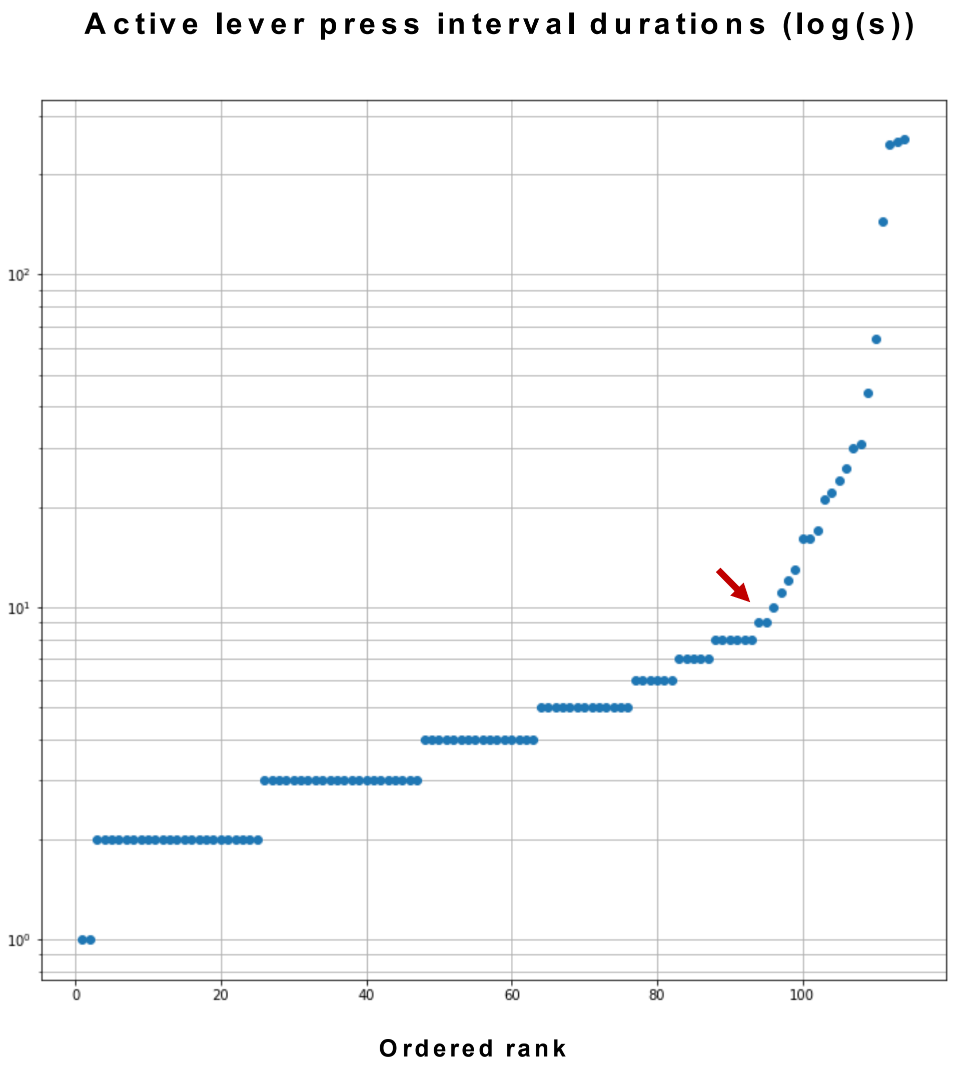


***Supplementary Figure 5.*** *Scree plot example with indication of inflection point estimation (red arrow).*

#### Supplemental Figure 6

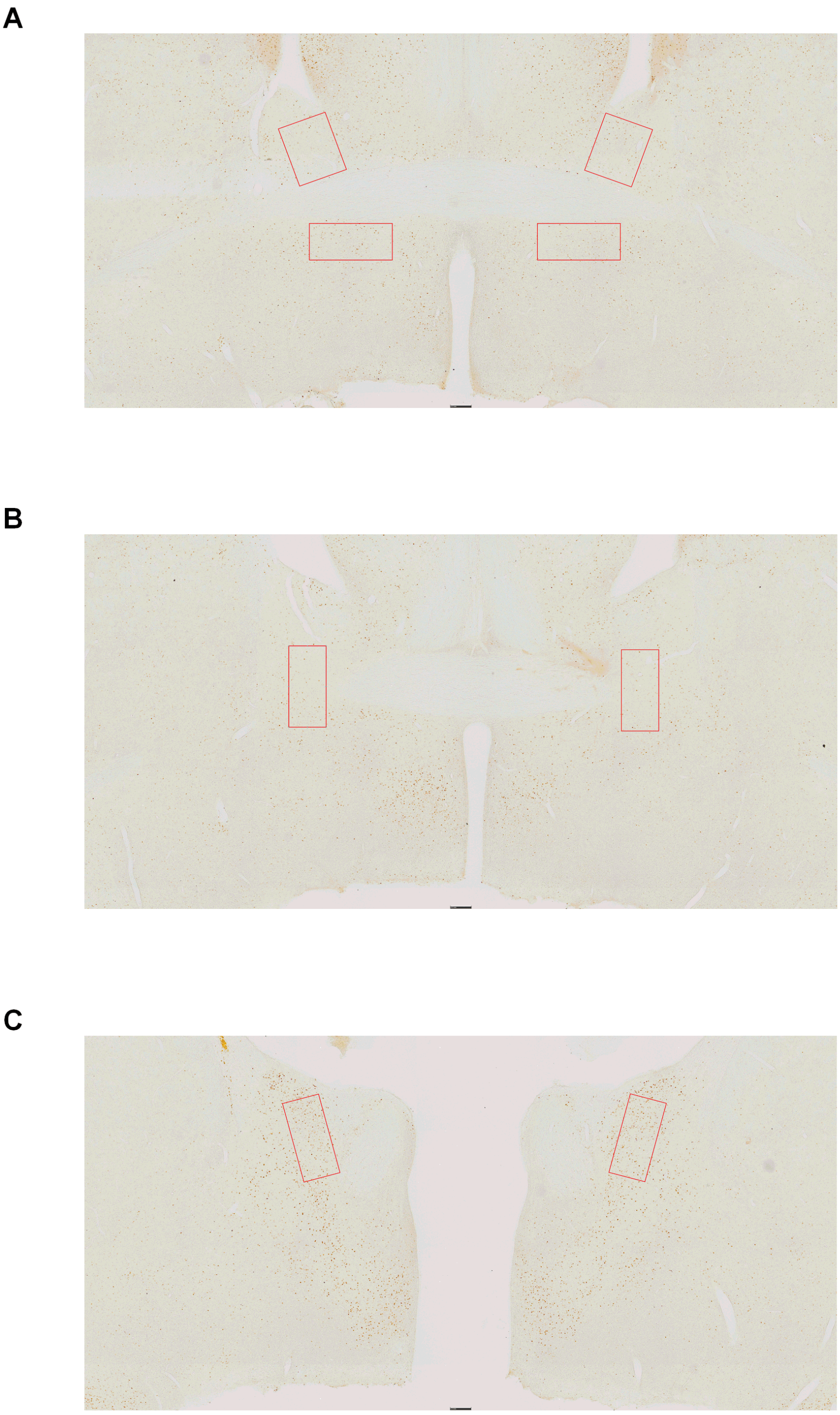

***Supplementary Figure 6.*** *Example images with ROI counting boxes locations on c-Fos stained sections. (A) Anterior (top boxes) and ventral (bottom boxes) BNST ROI's (AP between 0.0mm and -0.12mm). (B) Lateral BNST ROI's (AP between -0.24mm and -0.36mm). (C) Posterior BNST ROI's (AP between -0.72mm and -0.9mm).*

#### Supplemental Figure 7

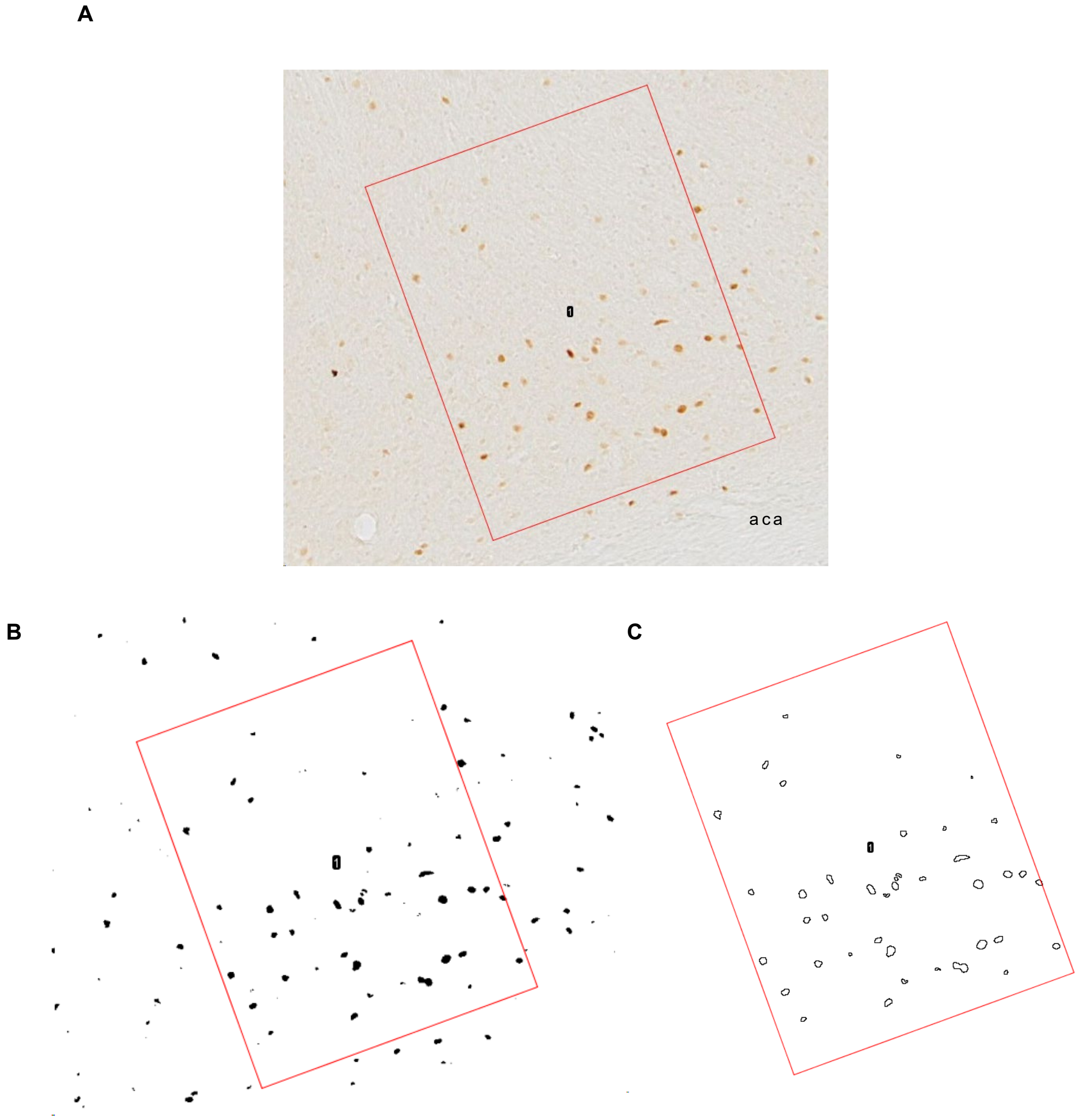


***Supplementary Figure 7*** *ImageJ c-Fos automated counting example. (A) Original image with a boxed area in the anterior BNST with c-Fos+ cells. aca; anterior commissure (B) Same image after thresholding and converting to binary. (C) Map of identified c-Fos+ cells by the particle analyze.*

**Table S1. Statistical analysis, related to Figures 1-4, S1-S3.**

| Figure | N per group | Statistical test | Test and p-values | Post hoc test |
|---|---|---|---|---|
| 1D | N=6 | One-tailed independent samples t-test | anterior t(10)=9.09, p<0.001; ventral t(10)=6.19, p<0.001 | |
| | | One tailed Mann-Whitney U test | lateral U=0, p=0.001; posterior U=0, p=0.001 | |
| 1E | Sham N=12<br>Gi N=11<br>Gq N=12 | Paired t-test - time in female vs male zone | VEH: Sham t(11)=6.19, p<0.001; Gi t(11)=6.83, p<0.001; Gq t(10)=10.46, p<0.001<br>CNO: Gi t(11)=5.58, p<0.001<br>Gq t(10)=3.69, p=0.004 | |
| | | Linear mixed model: virus*treatment | F(5, 63)= 4.69, p<0.001 | Gi: VEH-CNO p=0.007<br>Sham: VEH-CNO p=0.028 |
| 1F | Sham N=12<br>Gi N=11<br>Gq N=12 | One-sample t-test compared to 0.5 | VEH: Sham t(11)=6.34, p<0.001; Gi t(11)=9.24, p<0.001; Gq t(10)=12.76, p<0.001<br>CNO: Gi t(11)=6.29, p<0.001<br>Gq t(10)=3.84, p=0.003 | |
| | | Linear mixed model: virus*treatment | F(5, 63)= 4.69, p<0.001 | Gi: VEH-CNO p=0.054<br>Gq: VEH-CNO p=0.036 |
| 1G | Sham N=12<br>Gi N=11<br>Gq N=12 | Linear mixed model: virus*treatment | F(5,69)=7.12,p=0.023 | Gi: VEH-CNO p=0.023<br>CNO: Gi-Sham p=0.019 |
| 2B | Sham N=11 (left) and N=12 (right)<br>Gi N=12<br>Gq N=10 | Linear mixed model: virus*treatment | F(5,69)=4.26, p=0.002 | Gi: VEH-CNO p=0.035<br>CNO: Gi-Sham p=0.009 |
| 2C | Sham N=11 (left) and N=12 (right)<br>Gi N=12<br>Gq N=10 | Linear mixed model: virus*treatment | F(5,70)=3.04, p=0.015 | CNO: Gi-Sham p=0.020 |
| 2D | Sham N=11 (left) and N=12 (right)<br>Gi N=12<br>Gq N=10 | Linear mixed model: virus*treatment | NS | |
| 2E | Sham N=11 (left) and N=12 (right)<br>Gi N=12<br>Gq N=10 | Linear mixed model: virus*treatment | NS | |

| | | | | |
|---|---|---|---|---|
| 2F | Sham N=11 (left) and N=12 (right)<br>Gi N=12<br>Gq N=10 | Linear mixed model: virus*treatment | NS | |
| 2G | Sham N=11 (left) and N=12 (right)<br>Gi N=12<br>Gq N=10 | Linear mixed model: virus*treatment | F(5,66)=3.83, p<0.004 | Gi: VEH-CNO p=0.018 |
| 2H | Sham N=11 (left) and N=12 (right)<br>Gi N=12<br>Gq N=10 | Linear mixed model: virus*treatment | F(5,53)=2.79, p=0.026 | Gi: VEH-CNO p<0.001<br>CNO: Gi-Sham p<0.001 |
| 2I | Sham N=10 (left) and N=11 (right)<br>Gi N=11 (left) and N=8 (right)<br>Gq N=10 | Linear mixed model: virus*treatment | F(5,68)=3.58, p=0.006 | Gi: VEH-CNO p=0.002<br>CNO: Gi-Sham p=0.008 |
| 2J | Sham N=11 (left) and N=12 (right)<br>Gi N=12<br>Gq N=10 | Linear mixed model: virus*treatment | F(5,62)=8.96, p<0.001 | Gi: VEH-CNO p<0.001<br>CNO: Gi-Sham p<0.001 |
| 3B | HC, N=4,<br>FR1 N=5 | One-tailed independent samples t-test | Sham-FR1 vs. Sham-HC; anterior t(7)=2.46, p<0.022; lateral t(7)=2.90, p=0.012; posterior t(7)=2.48, p=0.021 | |
| 3C | Sham N=9<br>Gi N=11<br>Gq N=12 | Linear mixed model: virus*treatment | F(5,30)=4.14, p=0.006 | Gi: VEH-CNO p=0.005<br>CNO: Gi-Sham p=0.017<br>Gq-Sham p=0.023 |
| 3F | Sham N=9<br>Gi N=11<br>Gq N=12 | Linear mixed model: virus*treatment | NS | |
| 3G | Sham N=9<br>Gi N=11<br>Gq N=12 | Linear mixed model: virus*treatment | NS | |
| 3H | Sham N=9<br>Gi N=11<br>Gq N=12 | Linear mixed model: virus*treatment | F(5,33)=2.86, p=0.03 | Gi: VEH-CNO p=0.023 |
| 3I | Sham N=9<br>Gi N=11<br>Gq N=12 | Linear mixed model: virus*treatment | NS | |
| 3K | Sham N=9<br>Gi N=11 | Linear mixed model: virus*treatment | F(5,69)=3.18, p=0.012 | Gi: VEH-CNO p=0.006 |

| | | | | |
|---|---|---|---|---|
| | Gq N=12 | | | CNO: Gi-Sham p<0.001<br>Gq-Sham p=0.036 |
| 4A | Sham N=12<br>Gi N=12<br>Gq N=10 | Paired t-test - time in female vs male zone | VEH: Sham t(11)=7.98, p<0.001; Gi t(11)=9.07, p<0.001; Gq t(10)=6.95, p<0.001, and CNO: Sham t(11)=15.57, p<0.001; Gi t(11)=4.98, p<0.001; Gq t(10)=6.64, p<0.001. | |
| | | Linear mixed model (=1E):<br>- virus*treatment<br><br>- virus*treatment*condition | <br>F(5, 63)= 4.69, p<0.001<br><br>NS | <br>Gi: VEH-CNO p=0.012 |
| 4B | Sham N=12<br>Gi N=12<br>Gq N=10 | One-sample t-test compared to 0.5 | VEH: Sham t(11)=10.82, p<0.001; Gi t(11)=12.22, p<0.001; Gq t(10)=6.98, p<0.001, and CNO: Sham t(11)=29.37, p<0.001; Gi t(11)=5.14, p<0.001; Gq t(10)=7.01, p<0.001. | |
| | | Linear mixed model (=1F):<br>- virus*treatment<br><br>- virus*treatment*condition | <br>F(5, 68)= 3.53, p=0.007<br><br>NS | <br>Gi: VEH-CNO p=0.024 |
| 4C | Sham N=12<br>Gi N=12<br>Gq N=10 | Linear mixed model (=1G):<br>- virus*treatment<br><br><br>- virus*treatment*condition | <br>F(5,69)=7.12, p=0.023<br><br><br>NS | <br>Gi: VEH-CNO p<0.001<br>CNO: Gi-Sham p<0.001 |
| 4D | Sham N=12<br>Gi N=12<br>Gq N=10 | Linear mixed model (=2B):<br>- virus*treatment<br><br><br>- virus*treatment*condition | <br>F(5,69)=4.26, p=0.002<br><br><br>NS | <br>Gi: VEH-CNO p=0.006<br>CNO: Gi-Sham p=0.014 |
| 4E | Sham N=12<br>Gi N=12<br>Gq N=10 | Linear mixed model (=2C):<br>- virus*treatment<br><br>- virus*treatment*condition | <br>F(5,70)=3.04, p=0.015<br><br>NS | <br>Gi: VEH-CNO p=0.034 |
| 4F | Sham N=12<br>Gi N=12<br>Gq N=10 | Linear mixed model (=2H):<br>- virus*treatment<br>- virus*treatment*condition | <br>NS<br>NS | |
| 4G | Sham N=12<br>Gi N=12<br>Gq N=10 | Linear mixed model (=2I):<br>- virus*treatment<br><br>- virus*treatment*condition | <br>NS<br><br>F(5,67)=2.644, p=0.021 | <br><br><br>Pair-housing-Single housing: Gi-CNO p=0.002 |
| 4H | Sham N=12 | Linear mixed model (=2F): | | |

| | | | | |
|---|---|---|---|---|
| | Gi N=12<br>Gq N=10 | - virus*treatment<br><br>- virus*treatment*condition | F(5,67)=3.54, p=0.007<br><br>NS | Gi: VEH-CNO p=0.001 |
| 4I | Sham N=12<br>Gi N=12<br>Gq N=10 | Linear mixed model (=2G):<br>- virus*treatment<br><br><br>- virus*treatment*condition | <br>F(5,66)=3.83, p<0.004<br><br><br>NS | <br>Gi: VEH-CNO p=0.002<br>CNO: Gi-Sham p=0.044 |
| 4J | Sham N=11 (left) and N=12 (right)<br>Gi N=11 (left) and N=8 (right)<br>Gq N=18 (left) and N=11 (right) | Linear mixed model (=2J):<br>- virus*treatment<br><br><br>- virus*treatment*condition | <br>F(5,62)=8.96, p<0.001<br><br><br>NS | <br>Gi: VEH-CNO p=0.001<br>CNO: Gi-Sham p=0.006 |
| 4K | Sham N=8<br>Gi N=11<br>Gq N=12 | Linear mixed model (=3I):<br>- virus*treatment<br><br>- virus*treatment*condition | <br>NS<br><br>F(6,83)=24.90, p<0.001 | <br><br><br>*Ad libitum*-food restricted*:* p<0.001 across all groups (Sham, Gi, and Gq) and treatments) |
| 4L | Sham N=8<br>Gi N=11<br>Gq N=12 | Linear mixed model (=3K):<br>- virus*treatment<br><br>- virus*treatment*condition | <br>F(5,69)=3.18, p=0.012<br><br>F(6,84)=11.17, p<0.001 | <br>NS<br><br>*Ad libitum*-food restricted*:* Gi-CNO p<0.001, Gi-VEH p<0.001, Gq-CNO p<0.001, Gq-VEH p=0.020, Sham-CNO p=0.033, Sham-VEH p=0.016 |
| S1A | Sham N=6<br>Gq N=11 | One-tailed independent samples t-test<br><br><br>One tailed Mann-Whitney U test | anterior t(15)=1.95, p=0.035; ventral t(15)=2.03, p=0.030<br><br>lateral U=9, p=0.007; posterior U=0, p<0.001 | |
| S1B | Sham N=6<br>Gi N=11 | One-tailed Mann-Whitney U test | posterior U=15, p=0.026 | |
| S1C | Sham N=12<br>GI N=12<br>Gq N=11 | Linear mixed model: virus*treatment | NS | |
| S1D | Sham N=12<br>GI N=12<br>Gq N=11 | Linear mixed model: virus*treatment | NS | |

| | | | | |
|---|---|---|---|---|
| S1E | Sham N=4<br>Gq N=11 | One-tailed independent samples t-test<br><br>One-tailed Mann-Whitney U test | anterior t(13)=2.91, p=0.006<br><br>lateral U=0, p<0.001; posterior U=0, p<0.001 | |
| S1F | Sham N=5<br>Gi N=11 | One-tailed independent samples t-test or One-tailed Mann-Whitney U test | NS | |
| S2A | Pair housing:<br>Sham N=11<br>Gi N=12<br>Gq N=10<br>Single housing:<br>Sham N=12<br>Gi N=12<br>Gq N=11 | Linear mixed model:<br>- virus*treatment<br>- virus*treatment*condition | <br>NS<br>NS | |
| S2B | Pair housing:<br>Sham N=11<br>Gi N=12<br>Gq N=10<br>Single housing:<br>Sham N=12<br>Gi N=12<br>Gq N=11 | Linear mixed model:<br>- virus*treatment<br>- virus*treatment*condition | <br>NS<br>NS | |
| S2C | Pair housing:<br>Sham N=11<br>Gi N=12<br>Gq N=10<br>Single housing:<br>Sham N=12<br>Gi N=12<br>Gq N=11 | Linear mixed model:<br>- virus*treatment<br><br>- virus*treatment*condition | <br>F(5,71)=4.59, p=0.001<br><br>NS | Pair housing:<br>Gi: VEH-CNO: p=0.011<br>CNO: Gi-Sham: p=0.10<br>Single housing: post-hoc vs VEH: p=0.019; post-hoc vs Sham-CNO:p=0.005 |
| S2D | Pair housing:<br>Sham N=11<br>Gi N=12<br>Gq N=10<br>Single housing:<br>Sham N=12<br>Gi N=12<br>Gq N=11 | Linear mixed model:<br>- virus*treatment<br><br>- virus*treatment*condition | <br>F(5,62)=7.22, p<0.001<br><br>NS | Gi: VEH-CNO: p=0.002<br>CNO: Gi-Sham: p=0.001 |
| S2E | Pair housing: | Linear mixed model:<br>- virus*treatment | <br>F(5,68)=5.082, p=0.014 | <br>Single housing: |

| | | | | |
|---|---|---|---|---|
| | Sham N=11<br>Gi N=12<br>Gq N=10<br>Single housing:<br>Sham N=12<br>Gi N=12<br>Gq N=11 | - virus*treatment*condition | NS | Gq: VEH-CNO: p=0.003 |
| S2F | Pair housing:<br>Sham N=11<br>Gi N=12<br>Gq N=10<br>Single housing:<br>Sham N=12<br>Gi N=12<br>Gq N=11 | Linear mixed model:<br>- virus*treatment<br>- virus*treatment*condition | NS<br>NS | |
| S2G | Pair housing:<br>Sham N=11<br>Gi N=12<br>Gq N=10<br>Single housing:<br>Sham N=12<br>Gi N=12<br>Gq N=11 | Linear mixed model:<br>- virus*treatment<br>- virus*treatment*condition | F(5,55)=4.92, p<0.001<br>F(6,93)=2.74, p=0.017 | Gi: VEH-CNO: p<0.001<br>CNO: Gi-Sham: p=0.002<br>Pair-housing-Single-housing: Gi p<0.001<br>Gq p=0.004 |
| S2H | Pair housing:<br>Sham N=11<br>Gi N=12<br>Gq N=10<br>Single housing:<br>Sham N=12<br>Gi N=12<br>Gq N=11 | Linear mixed model:<br>- virus*treatment<br>- virus*treatment*condition | F(5,65)=6.92, p<0.001<br>NS | Pair-housing:<br>Gi: VEH-CNO: p=0.001<br>CNO: Gi-Sham: p=0.013<br>Single housing:<br>Gi: VEH-CNO p=0.002<br>CNO: Gi-sham p=0.006 |
| S3A | Sham N=9<br>GI N=11<br>Gq N=12 | Linear mixed model:<br>virus*treatment | NS | |
| S3B | Ad libitum:<br>Sham N=9<br>Gi N=11<br>Gq N=12<br>Food restriction:<br>Sham N=8<br>Gi N=11<br>Gq N=12 | Linear mixed model:<br>- virus*treatment<br>- virus*treatment*condition | NS<br>F(6,84)=14.72, p<0.001 | *Ad libitum*-food restricted: Gi-CNO p=0.001; Gi-VEH p=0.002; p<0.001 across the other |

| | | | | |
|---|---|---|---|---|
| | | | | groups and treatments |
| S3C | Ad libitum:<br>Sham N=9<br>Gi N=11<br>Gq N=12<br>Food restriction:<br>Sham N=8<br>Gi N=11<br>Gq N=12 | Linear mixed model:<br>- virus*treatment<br><br>- virus*treatment*condition | <br>NS<br><br>F(6,84)=17.240, p<0.001 | <br><br><br>*Ad libitum*-food restricted: Gi-VEH p=0.001; p<0.001 across the other groups and treatments |
| S3D | Ad libitum:<br>Sham N=9<br>Gi N=11<br>Gq N=12<br>Food restriction:<br>Sham N=8<br>Gi N=11<br>Gq N=12 | Linear mixed model:<br>- virus*treatment<br>- virus*treatment*condition | <br>NS<br>NS | |
| S3E | Ad libitum:<br>Sham N=9<br>Gi N=11<br>Gq N=12<br>Food restriction:<br>Sham N=8<br>Gi N=11<br>Gq N=12 | Linear mixed model:<br>- virus*treatment<br><br>- virus*treatment*condition | <br>F(5,68)=4.41, p=0.002<br><br>F(6,79)=2.62; p=0.046 | <br>*Ad libitum:* Gq: VEH-CNO: p=0.002<br>CNO: Gq-Sham: p=0.026<br>Food restricted: Gq: VEH-CNO: p=0.019<br><br>*Ad libitum*-food restricted:<br>Sham-VEH p=0.040 |
